# The effects of rapid mitochondrial gene loss on organellar proteomes

**DOI:** 10.1101/2025.11.24.690252

**Authors:** Jessica M. Warren, Amanda K. Broz, Ryan Stikeleather, Daniel B. Sloan

## Abstract

Mitochondrial genomes retain only a tiny number of genes from their bacterial progenitors, including key components of protein translation machinery. The set of mitochondrially encoded tRNAs and ribosomal subunits is highly variable across angiosperms, with many examples of mitochondrial gene loss, replacement, and/or transfer to the nucleus. This dynamic history suggests large-scale remodeling of mitochondrial translation machinery in some lineages, but such conclusions are largely inferred from genomic sequence and protein targeting predictions. Here, we use proteomic (LC-MS/MS) analysis of purified mitochondria and chloroplasts from angiosperm species with major differences in mitochondrial gene content (*Arabidopsis thaliana* and *Silene conica*). Our analysis largely confirms the current understanding of subcellular localization for nuclear-encoded proteins involved in tRNA metabolism and ribosome function in *A. thaliana*, although some aminoacyl-tRNA synthetases (aaRSs) may have more specialized subcellular roles than previously thought. In contrast, *S. conica* has undergone extensive mitochondrial gene loss and numerous associated changes in the composition of its mitochondrial proteome, including apparent retargeting of aaRSs, replacement of ribosomal subunits, and loss of the glutamine amidotransferase (GatCAB) complex. Overall, this analysis illustrates how the complex network of molecular interactions necessary for mitochondrial translation are perturbed by gene loss, transfer, and replacement.

**Significance Statement:** Plant mitochondrial genomes exhibit exceptional diversity in gene content across species due to ongoing processes of gene loss, transfer, and functional replacement. However, our understanding of the consequences of these genomic changes on the mitochondrial proteome remain limited. Here, we compare organellar (mitochondrial and chloroplast) proteomes across species of flowering plants, revealing a large-scale rewiring of mitochondrial protein synthesis machinery has coincided with loss of tRNA and ribosomal protein genes from the mitochondrial genome.

## Introduction

Endosymbiotically derived organelles such as mitochondria and plastids evolved from free-living bacteria and still retain their own genomes after more than a billion years (Gould et al. 2008; Roger et al. 2017). Thus, protein synthesis occurs within each of these organelles, using machinery that is distinct from the translation system responsible for synthesizing nuclear-encoded proteins in the cytosol. One common theme in the evolution of obligate endosymbionts is extensive gene loss, functional replacement, and/or transfer to the nucleus (McCutcheon and Moran 2012; Sloan et al. 2018). As such, the gene content in ancient endosymbionts/organelles is whittled down to mostly just key components of the metabolic and biosynthetic functions they provide to the host cell, such as cellular respiration in mitochondria and photosynthesis in plastids. However, even the most ancient endosymbionts/organelles retain some of the genes involved in translation, including ribosomal rRNAs (rRNAs), ribosomal protein subunits, and/or transfer RNAs (tRNAs) (Timmis et al. 2004; Salinas-Giegé et al. 2015; McCutcheon et al. 2024).

Mitochondrial gene content has largely stabilized in some eukaryotic lineages. For example, most bilaterian animals retain the same set of 37 genes that were ancestrally present prior to the Cambrian Explosion (Boore 1999). In contrast, mitochondrial gene content is highly dynamic in angiosperms, often differing even among closely related species (Adams, Qiu, et al. 2002). Angiosperm mitochondrial genomes (mitogenomes) can contain anywhere from 19 to 41 protein-coding genes and from 1 to 19 types of tRNA genes (excluding gene duplicates) (Richardson et al. 2013; Skippington et al. 2015; Yu et al. 2025).

Multiple evolutionary processes can facilitate the loss of genes from the mitogenome. Recent losses of mitochondrial protein-coding genes in angiosperms are typically associated with transfer of those genes to the nucleus. Prior to the loss of the native mitochondrial gene copy, the transferred nuclear copy must be expressed, and the resulting protein must be imported back into the mitochondria (Adams et al. 1999; Sloan et al. 2018). In addition to mitochondrial-to-nuclear gene transfer, there are also cases where mitochondrial protein-coding genes are functionally replaced by homologs of plastid or nuclear origin (Adams, Daley, et al. 2002). Losses of mitochondrial tRNA genes appear to follow this latter route. Specifically, they are replaced by import of existing nuclear-encoded tRNAs from the cytosol (Salinas-Giegé et al. 2015; Warren et al. 2021). To our knowledge, there are no documented cases in which a mitochondrial tRNA gene was functionally transferred to the nucleus and targeted back for import into the mitochondria (although likely cases of plastid tRNA gene transfer were recently discovered in the lycophyte *Selaginella* and parasitic angiosperms in the family Balanophoraceae; Berrissou et al. 2024; Ceriotti et al. 2026).

A fundamental challenge in the field of cytonuclear coevolution is to understand the process by which functional gene replacements occur and whether they perturb the intimate and coevolved interactions among gene products encoded in two different genomes. For example, tRNAs must be recognized by their cognate aminoacyl-tRNA synthetases (aaRSs) to be charged with the correct amino acid, and ribosomal proteins and rRNAs physically interact with dozens of subunits within a massive enzyme complex. It is remarkable that functional replacement of a (bacterial-like) mitochondrial gene with its (archaeal-like) nuclear counterpart is possible given that their divergence spans the very deepest split in the tree of life and that even small sequence changes have the potential to disrupt these coevolved interactions (Meiklejohn et al. 2013; Sloan et al. 2023).

The mitogenomes found in the angiosperm genus *Silene* are highly variable among species and characterized by many unusual features. Perhaps the most extreme of these mitogenomes is found in *S. conica*. It has a highly accelerated mutation rate, the largest size of any known angiosperm mitogenome (>11 Mb), and a fragmented multichromosomal structure (Sloan, Alverson, Chuckalovcak, et al. 2012; Broz et al. 2021). Despite its large size, the *S. conica* mitogenome has an unusually small gene content with just 25 protein-coding, 3 rRNA, and 2 tRNA genes (excluding gene duplicates). The near-complete loss of tRNA genes from this mitogenome has been accompanied by extensive import of cytosolic-like tRNA counterparts (Warren et al. 2021). Our previous analysis based on *in silico* targeting predictions and fluorescence microscopy indicated that these tRNA replacement events in *S. conica* have had differing effects on coevolved relationships with aaRSs (Warren et al. 2023). All plant aaRSs are nuclear-encoded, but they differ in where they are localized within the cell. One of the most common patterns is that the plant expresses two aaRSs for a given amino acid – one that functions in the cytosol and another that is dual-targeted and imported into both the mitochondria and plastids (Duchêne et al. 2005). For half of the replaced tRNA genes in *S. conica*, we found evidence that a corresponding cytosolic aaRS also gained targeting to the mitochondria. Therefore, the ancestral pairing of a cytosolic tRNA and aaRS was apparently maintained in these cases and simply relocated to an additional cellular compartment. In contrast, we did not find evidence of aaRS retargeting for the other half of replaced tRNA genes, suggesting that (bacterial-like) organellar aaRSs are responsible for charging the cytosolic-like tRNAs now being imported into the mitochondria. However, these inferences have not been investigated by direct analysis of mitochondrial protein content, as the mitochondrial proteome of *S. conica* remains entirely unexplored.

Here, we perform proteomic analysis of *S. conica* and the model angiosperm *Arabidopsis thaliana* using liquid chromatography-tandem mass spectrometry (LC-MS/MS). The resulting datasets allow us to directly investigate how the composition of the mitochondrial proteome and its translational machinery have changed in association with the extensive gene loss from the *S. conica* mitogenome.

## Results and Discussion

### Purification of mitochondrial and chloroplast proteomes

We analyzed LC-MS/MS data from extracted protein content from two biological replicates of purified mitochondria, purified chloroplasts, and total leaf tissue from both *A. thaliana* and *S. conica*. A similar dataset was also generated for a third species (*Agrostemma githago*). However, preliminary analysis of the *A. githago* dataset indicated that it had limited detection of the translation machinery that was the focus of this study. Therefore, we have deposited data from all three species (see Data Availability), but only *A. thaliana* and *S. conica* were analyzed in the present study.

We used two (semi-)quantitative metrics of protein abundance: peptide-spectrum matches (PSMs) and normalized MS1 ion intensity (i.e. peak area). Overall, we detected PSMs from 5599 and 5491different proteins in *A. thaliana* and *S. conica*, respectively (Table 1). To verify the efficacy of our organelle purifications, we measured the enrichment of each mitochondrial-encoded and plastid-encoded protein in the purified organelles relative to total leaf tissue. We used these proteins because they are not expected to be exported to other cellular compartments and, therefore, represent the most reliable available markers for their respective organelles. Although we did detect signal from both ion intensity (Figure 1) and PSMs (Figure S1) for mitochondrial-encoded proteins in chloroplast samples and vice versa, we observed a strong quantitative difference between the sample types that confirmed we had effectively enriched for mitochondrial and chloroplast subcellular fractions. Our purified mitochondrial fractions from *A. thaliana* exhibited an average enrichment in cumulative mitochondrial-encoded peptide ion intensity of 18-fold relative to total leaf tissue and 554-fold relative to purified chloroplasts. Likewise, *S. conica* mitochondrial fractions showed 15-fold enrichment relative to leaf tissue and 157-fold relative to purified chloroplasts (Figure 1A). These patterns were mirrored by separation of the sample types at the level of individual proteins (Figure 1B). We also saw clear separation among sample types based on PSM counts. Enrichment ratios calculated based on PSMs tended to be lower (Figure S1), which was expected because “dynamic exclusion” settings during LC-MS/MS data-dependent acquisition lead to intentional downsampling of highly abundant PSMs to increase overall peptide detection.

**Figure 1.**
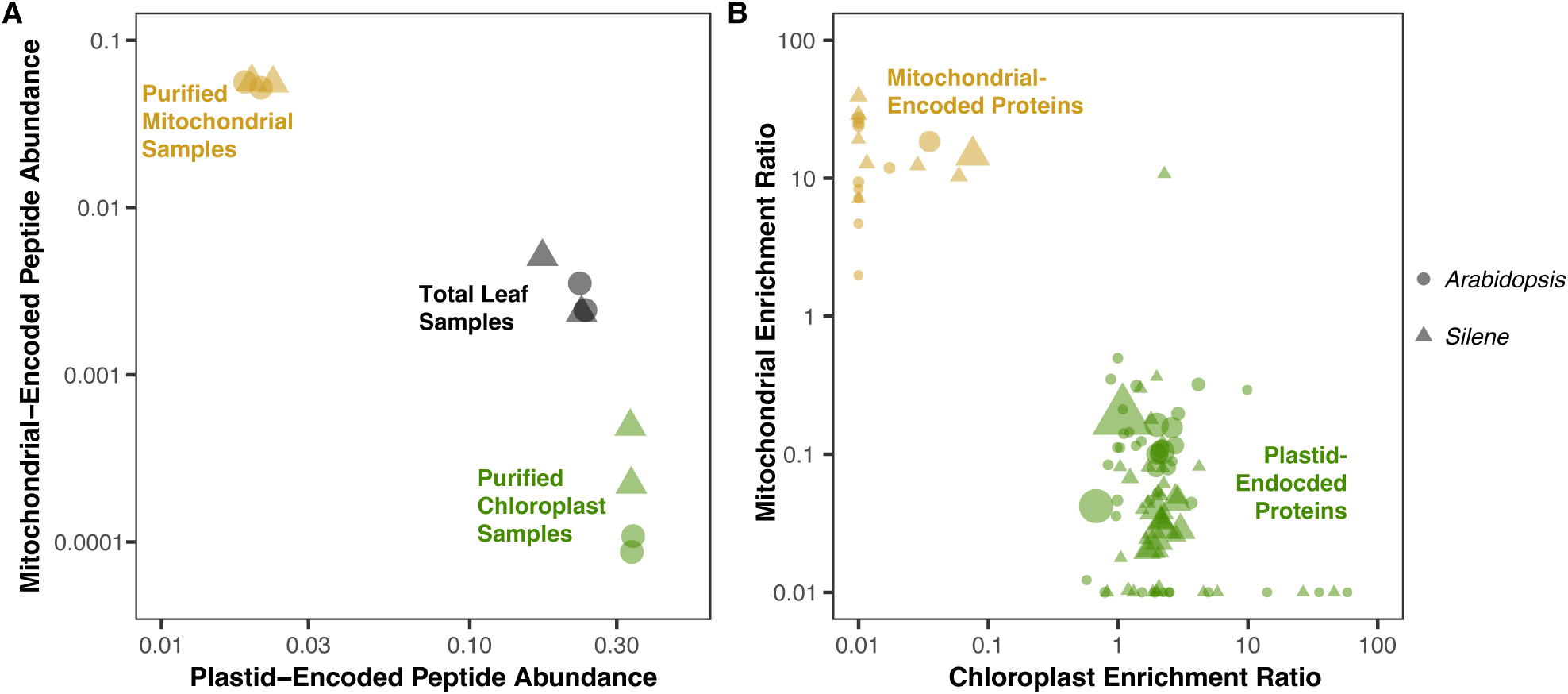
Enrichment of proteins encoded by the mitochondrial and plastid genomes in purified organelle fractions. (A) Each point in this panel represents an individual biological sample, showing its cumulative ion intensity across all plastid-encoded proteins (x-axis) and mitochondrial-encoded proteins (y-axis) expressed as a proportion of all protein abundance in the sample with each biological replicate shown separately. (B) Each point in this panel represents a mitochondrial-encoded or plastid-encoded protein. Enrichment in chloroplast samples (x-axis) or mitochondrial samples (y-axis) is calculated by dividing the ion intensity for that protein from the respective purified organelles by the corresponding ion intensity from total leaf samples. Point size is scaled based on total abundance of the protein across all samples. Only proteins represented by at least 5 unique peptides in the dataset are shown, and enrichment/depletion values were capped at 100-fold for visualization purposes. The two biological replicates are averaged for this panel. In both panels, point shape indicates species identity (circle: *A. thaliana*; triangle: *S. conica*), and ion intensities were excluded for peptides that were shared between multiple proteins or if they were identified solely based on an MS1 peak that was not validated in that sample with an MS2 spectrum. An equivalent analysis based on PSMs is available in Figure S1.

**Table 1.**
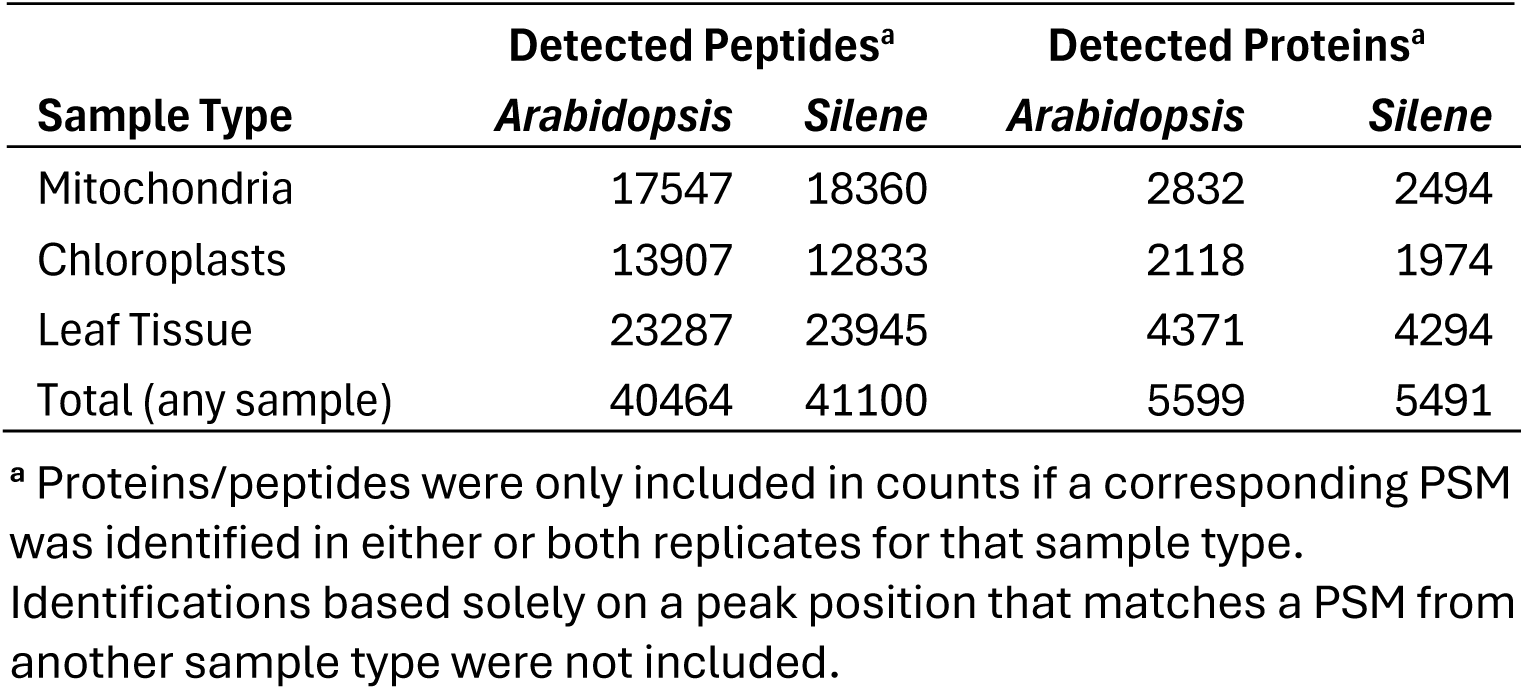
Detected proteins in LC-MS/MS dataset.

| **Sample Type** | **Detected Peptides<sup>a</sup>** |  | **Detected Proteins<sup>a</sup>** |  |
| --- | --- | --- | --- | --- |
|  | ***Arabidopsis*** | ***Silene*** | ***Arabidopsis*** | ***Silene*** |
| Mitochondria | 17547 | 18360 | 2832 | 2494 |
| Chloroplasts | 13907 | 12833 | 2118 | 1974 |
| Leaf Tissue | 23287 | 23945 | 4371 | 4294 |
| Total (any sample) | 40464 | 41100 | 5599 | 5491 |

<sup>a</sup> Proteins/peptides were only included in counts if a corresponding PSM was identified in either or both replicates for that sample type. Identifications based solely on a peak position that matches a PSM from another sample type were not included.

In *A. thaliana*, we also made use of the SUBA5 consensus classification (August 2022 release) to assess the effectiveness of our mitochondrial and chloroplast purifications (Hooper et al. 2014). Of the 5599 proteins we detected in *A. thaliana*, SUBA5 identifies 980 and 1477 of them as targeted to mitochondria and plastids, respectively. As expected, the purified mitochondrial and chloroplast samples showed large and opposite shifts in composition with respect to these proteins (Table S1).

We also performed correlation analyses among samples within each species to test for consistency between pairs of biological replicates. As expected, replicate mitochondrial samples and replicate chloroplast samples all exhibited high correlation coefficients (0.78 to 0.93; Figures S2-S5). In contrast, mitochondrial samples had low correlation coefficients (≤ 0.33) with the other two sample types. Chloroplast and total leaf samples showed intermediate levels of correlation with each other (Figures S2-S5). This correlation likely reflects the fact that chloroplasts account for the majority of the total proteome in leaf cells (Heinemann et al. 2021), which also sets a limit to the maximum proportional enrichment that can be obtained for chloroplast proteins relative to total leaf tissue.

### Subcellular specialization of Arabidopsis aaRSs

Numerous studies have been conducted to catalog the subcellular localization of aaRSs in *A. thaliana*, primarily by fusing putative organelle-targeting transit peptides from aaRSs to fluorescent proteins or other reporters, which could be visualized within cells or tested for import into isolated organelles *in vitro* (Mireau et al. 1996; Uwer et al. 1998; Souciet et al. 1999; Peeters et al. 2000; Duchêne et al. 2001; Duchêne et al. 2005). Collectively, these studies established a consensus for the subcellular localization of all aaRSs in *A. thaliana* (Table 2), which we will refer to as the Duchêne classification because the most extensive sampling was performed by Duchêne et al. (2005), and the cumulative body of work was later summarized by Duchêne et al. (2009). For most amino acids, *A. thaliana* expresses two different aaRSs – one that is targeted to the cytosol (cyto-only) and another that is dual-targeted to both the chloroplasts and mitochondria (chloro-mito). However, some aaRSs exhibit atypical patterns of subcellular localization, such as dual targeting to both the cytosol and mitochondria (cyto-mito) or to all three cellular compartments. One LeuRS enzyme (AT4G04350) was also found to be specific to the chloroplasts (chloro-only; Duchêne et al. 2009).

**Table 2.** Comparison between aaRS detection in LC-MS/MS datasets from this study and others (Fuchs et al. 2020; van Wijk et al. 2021; Rugen et al. 2024) and previous classification of subcellular localization (Duchêne et al. 2005; Duchêne et al. 2009)

| Accession | Type | Duchêne Classification | Notes on Potential Refinements<sup>c</sup> |
| --- | --- | --- | --- |
| AT5G22800 | AlaRS | Chloro-Mito | No mito localization detected; may effectively function as chloro-only |
| AT1G50200 | AlaRS | Cyto-Chloro-Mito | No chloroplast localization detected; may effectively function as cyto-mito |
| AT4G26300 | ArgRS | Cyto-Chloro-Mito(?) | Mito localization confirmed |
| AT1G66530 | ArgRS | Cyto-only |  |
| AT4G17300 | AsnRS | Chloro-Mito |  |
| AT5G56680 | AsnRS | Cyto-only |  |
| AT1G70980<sup>a</sup> | AsnRS | Cyto-only |  |
| AT4G33760 | AspRS | Chloro-Mito |  |
| AT4G31180 | AspRS | Cyto-only |  |
| AT4G26870 | AspRS | Cyto-only |  |
| AT2G31170 | CysRS | Chloro-Mito |  |
| AT5G38830 | CysRS | Cyto-only |  |
| AT3G56300<sup>a</sup> | CysRS | Cyto-only |  |
| AT1G25350 | GlnRS | Cyto-only |  |
| AT5G64050 | GluRS | Chloro-Mito |  |
| AT5G26710 | GluRS | Cyto-only |  |
| AT3G48110 | GlyRS | Chloro-Mito |  |
| AT1G29880 | GlyRS | Cyto-Mito<sup>b</sup> |  |
| AT3G46100 | HisRS | Chloro-Mito |  |
| AT3G02760 | HisRS | Cyto-only |  |
| AT5G49030 | IleRS | Chloro-Mito |  |
| AT4G10320 | IleRS | Cyto-only |  |
| AT4G04350 | LeuRS | Chloro-only |  |
| AT1G09620 | LeuRS | Cyto-Mito |  |
| AT3G13490 | LysRS | Chloro-Mito |  |
| AT3G11710 | LysRS | Cyto-only |  |
| AT3G55400 | MetRS | Chloro-Mito |  |
| AT4G13780 | MetRS | Cyto-only |  |
| AT3G58140 | PheRS | Chloro-Mito |  |
| AT4G39280 | PheRS | Cyto-only |  |
| AT1G72550 | PheRS | Cyto-only |  |
| AT5G52520 | ProRS | Chloro-Mito |  |
| AT3G62120 | ProRS | Cyto-only |  |
| AT1G11870 | SerRS | Chloro-Mito |  |
| AT5G27470 | SerRS | Cyto-only |  |
| AT2G04842 | ThrRS | Chloro-Mito | No mito localization detected; may effectively function as chloro-only |
| AT5G26830 | ThrRS | Cyto-Mito |  |
| AT1G17960<sup>a</sup> | ThrRS | Cyto-only |  |
| AT2G25840 | TrpRS | Chloro-Mito |  |
| AT3G04600 | TrpRS | Cyto-only |  |
| AT3G02660 | TyrRS | Chloro-Mito |  |
| AT1G28350 | TyrRS | Cyto-only |  |
| AT2G33840<sup>a</sup> | TyrRS | Cyto-only |  |
| AT5G16715 | ValRS | Chloro-Mito | Low mito localization detected; may effectively function as chloro-only |
| AT1G14610 | ValRS | Cyto-Mito<sup>b</sup> | May be functional in mito given low detection of AT5G16715 in mito |

<sup>a</sup> Four aaRSs were not detected in our proteomic dataset (all predicted to be cyto-only and close paralogs of aaRSs that were detected).
<sup>b</sup> The cyto-mito GlyRS and ValRS enzymes have previously suggested to not function in mitochondrial aminoacylation despite localization to the organelle (Duchêne et al. 2001; Duchêne et al. 2009).
<sup>c</sup> References to no detected mitochondrial/chloroplast localization are based on the absence of any PSMs from our samples. Other studies have detected these proteins at low levels (see main text and Figure 3).

Overall, our patterns of aaRS enrichment in mitochondrial and chloroplast fractions strongly aligned with the Duchêne classification (Figure 2; Table 2). All aaRSs that had been previously described as cyto-only showed little or no detection in our purified mitochondrial and chloroplasts samples from *A. thaliana*. In contrast, all of these aaRSs were detected in total leaf tissue (except for four that presumably had low overall expression or detectability due to the presence of close paralogs with higher expression; Table 2), supporting the inference that their functional role is limited to the cytosol. Likewise, the four previously identified cyto-mito aaRSs (GlyRS, LeuRS, ValRS, and ThrRS) were found at higher abundance in mitochondria than all the cyto-only aaRSs and were not detected in chloroplasts. As expected, previously identified chloro-mito aaRSs were generally detected at high abundance in both organelle fractions, and the chloro-only LeuRS was found in high abundance in our chloroplast fractions but not detected in mitochondria (Figure 2; Table 2). In addition, our data support previous inferences that one ArgRS enzyme (AT4G26300) functions in all three cellular compartments. In prior GFP localization studies, this ArgRS protein was only observed to be targeted to the chloroplasts (Duchêne et al. 2005), but it was predicted to function in the cytosol and mitochondria as well because mutants lacking the only other known *A. thaliana* ArgRS gene (AT1G66530) are still viable (Berg et al. 2005; Duchêne et al. 2009). Indeed, we detected AT4G26300 in both mitochondrial and chloroplast fractions (Figure 2). In contrast, AT1G66530 was only detected in total leaf samples and at very low abundance (only a single PSM in each replicate sample). Therefore, it is likely that both ArgRS enzymes are expressed in the cytosol, but whether AT1G66530 makes any contribution to cytosolic translation is not clear.

**Figure 2.**
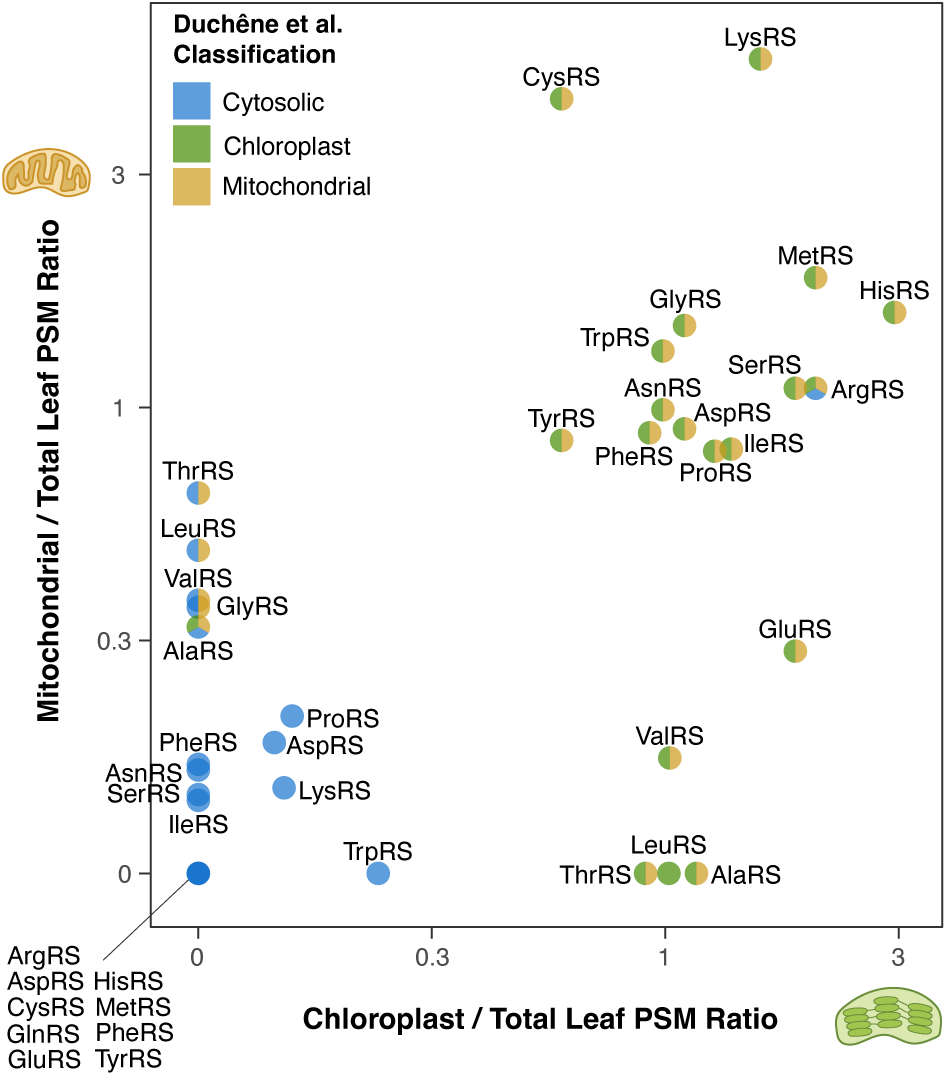
Summary of mitochondrial and chloroplast enrichment of *Arabidopsis thaliana* aaRSs relative to total leaf samples based on ratios of PSM counts combined across two biological replicates. Color coding of points reflects whether the aaRS was previously classified as being targeted to the cytosol, chloroplasts, and/or mitochondria (Duchêne et al. 2005; Duchêne et al. 2009). A similar analysis based on ion intensity is available in Figure S6.

Despite the overall congruence between our proteomic analysis and the Duchêne classification, there were some discrepancies that point to potential refinements in our current understanding of aaRS targeting in *A. thaliana* (Table 2). First, a previous analysis has indicated that there is one chloro-mito AlaRS (AT5G22800) and another AlaRS (AT1G50200) targeted to all three cellular compartments (Duchêne et al. 2005). However, we did not detect AT1G50200 in purified chloroplasts, and we did not detect AT5G22800 in purified mitochondria (Figure 2), implying that they exist at low abundances in these organelles.

Therefore, it is possible that AlaRSs follow a division of labor similar to the pattern observed for LeuRSs, with effectively distinct cyto-mito and chloro-only enzymes. This scenario would be consistent with the finding that plastid tRNA-Ala is a poor substrate for the cytosolic AlaRS in spinach (Steinmetz and Weil 1986) and that the fusion of the AT1G50200 transit peptide to β-glucuronidase (GUS) failed to localize reporter activity to chloroplasts (Mireau et al. 1996). However, the lines of evidence reported by Duchêne et al. (2005) supporting dual-organellar localization of both AlaRSs suggest that AT1G50200 and AT5G22800 are indeed present in chloroplasts and mitochondria, respectively, but likely at low levels we were unable to detect with our analysis. Accordingly, some other proteomic studies have detected AT1G50200 at low levels in *A. thaliana* chloroplasts (van Wijk et al. 2021), and as we report below, mining other mitochondrial proteomic datasets from *A. thaliana* also supports this conclusion.

Second, the ThrRS enzymes might offer yet another example of an effective division of labor between cyto-mito and chloro-only enzymes in *A. thaliana*, even though previous analysis has suggested two different ThrRSs functioning in the mitochondria (one chloro-mito and one cyto-mito). Fusion of the transit peptide from the putative chloro-mito ThrRS (AT2G04842) to reporter genes was previously shown to drive localization and import into both chloroplasts and mitochondria (Duchêne et al. 2005), but we only detected this ThrRS in chloroplasts. Instead, the most abundant ThrRS in mitochondria was the previously classified cyto-mito ThrRS (AT5G26830), and as expected, we did not detect this ThrRS in chloroplasts. A third ThrRS gene (AT1G17960) has been identified in *A. thaliana* and presumed to have cyto-only function due to the apparent lack of any transit peptide (Duchêne et al. 2005), but we did not detect this protein in any of our samples.

Third, our results may provide insights into the unusual cases of GlyRS and ValRS function in *A. thaliana*. Although there is a cyto-mito and a chloro-mito aaRS reported in each case, the cyto-mito GlyRS has been suggested to not function in aminoacylation within the mitochondria (Duchêne et al. 2001). Likewise, the cyto-mito ValRS might not be active in mitochondrial aminoacylation according to a personal communication reported by Duchêne et al. (2009). We detected the chloro-mito GlyRS (AT3G48110) at substantial abundance in mitochondria (Figure 2), as expected if it truly is the sole or primary enzyme responsible for aminoacylation of mitochondrial tRNA-Gly even though the cyto-mito GlyRS (AT1G29880) is also present in the mitochondrial fraction. In contrast, we detected the putative chloro-mito ValRS (AT5G16715) at only very low abundance in *A. thaliana* mitochondria (only a single PSM in one of the two replicate samples; Figure 2). Coupled with the fact that there has been conflicting or inconsistent evidence for targeting of AT5G16715 to mitochondria (Duchêne et al. 2005; Duchêne et al. 2009), this observation raises the possibility that ValRS represents a fourth example (along with AlaRS, LeuRS, and ThrRS) where the division of labor is primarily between a cyto-mito and a chloro-only enzyme.

To further investigate the degree of aaRS subcellular specialization, we analyzed data from two previously published studies that obtained deeper and more quantitative coverage of the *A. thaliana* mitochondrial proteome than our own analysis (Fuchs et al. 2020; Rugen et al. 2024). Both studies detected nearly every aaRS in the mitochondrial proteome (Figure 3A), including ones that have been characterized as cyto-only. The fact that these cyto-only aaRSs were generally detected at low abundances makes it difficult to interpret whether there truly is ubiquitous (albeit low-level) import of these enzymes into mitochondria or whether some detection simply reflects small amounts of contamination in purified mitochondrial fraction. Regardless, these studies provide high-quality quantitative estimates that are in good agreement with our observations. Data from both studies support the conclusion that the three aaRSs discussed above (AlaRS, ThrRS, and ValRS) have the lowest mitochondrial abundance of any of the previously classified mito-chloro aaRSs. Indeed, some estimates from Fuchs et al. (2020) for these three were lower than many aaRSs that have been previously classified as cyto-only (Figure 3A). Overall, mitochondrial abundances for alternative aaRSs from the same amino-acid family exhibit a strong negative correlation that is spread across a wide continuum (Figure 3B).

**Figure 3.**
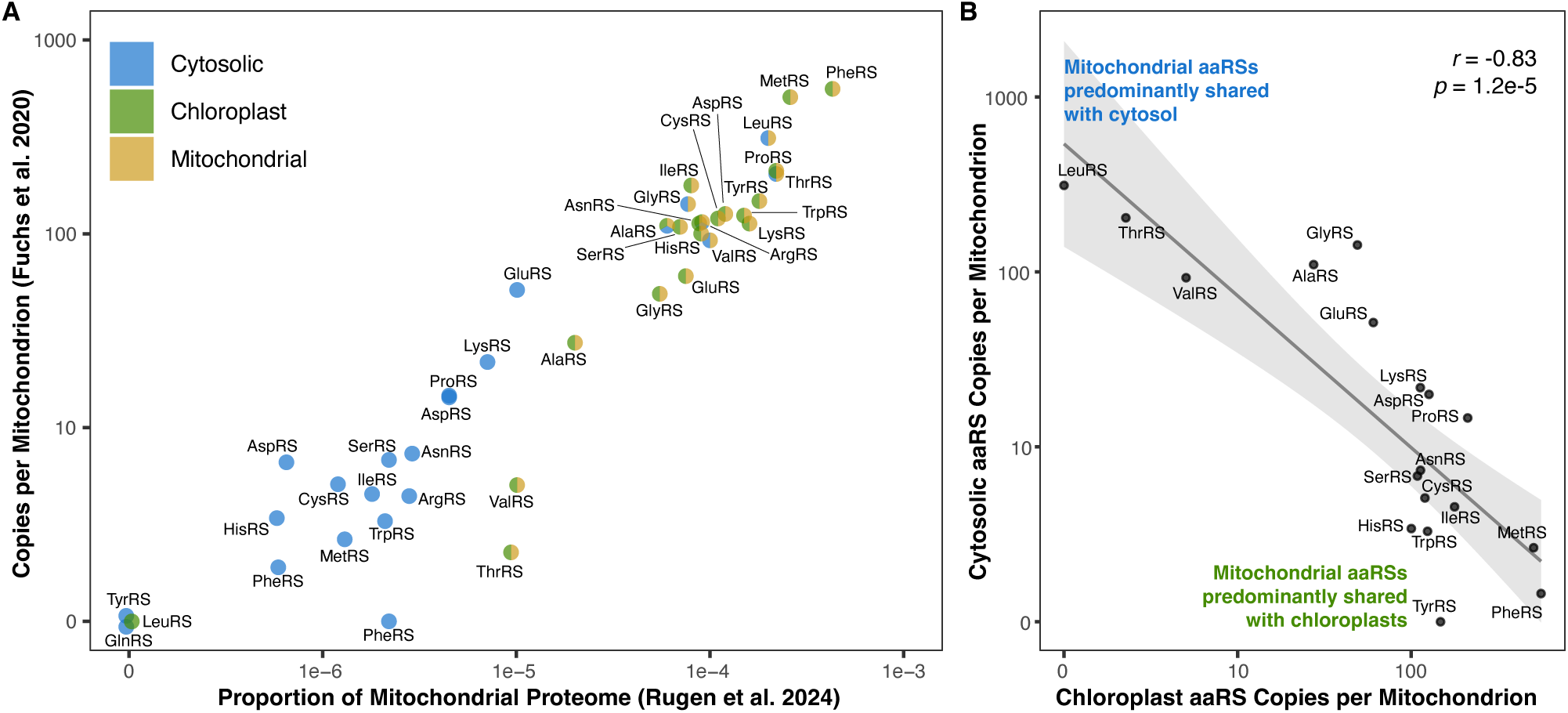
Summary of aaRS abundance from published *A. thaliana* mitochondrial proteome studies. (A) Quantifications from two previous LC-MS/MS datasets (Fuchs et al. 2020; Rugen et al. 2024) generally show strong association between measured abundance in mitochondria and whether the enzyme was previously classified as mitochondrial-targeted as indicated by color coding of points reflects whether the aaRS was previously classified as being targeted to the cytosol, chloroplasts, and/or mitochondria (Duchêne et al. 2005; Duchêne et al. 2009). However, the AlaRS, ThrRS, and ValRS enzymes classified as chloro-mito show very low mitochondrial findings consistent with our findings (Figure 2). Note that the chloro-only LeuRS and the cyto-only GlnRS and TyrRS were not detected in either study, but their points are slightly offset in the plot for visibility. (B) aaRS families exhibit a negative correlation in mitochondrial abundance between enzymes that are shared with the cytosol and those that are shared with the chloroplasts. Each point represents an aaRS family, where the x- and y-values indicate the mitochondrial abundances from Fuchs et al. (2020) for members of that family that are (also) found in the chloroplasts or cytosol, respectively. ArgRS is excluded from this plot because the highly expressed enzyme (AT4G26300) in this family is shared between all three compartments, and GlnRS is excluded because there is only a single enzyme (AT1G25350) in the family due to the use of the indirect tRNA-Gln aminoacylation pathway in mitochondria and plastids (see main text). The AlaRS localized to all three compartments (AT1G50200) was treated as cytosolic for the purposes of this analysis due to its low abundance in the chloroplasts (Figure 2).

Therefore, these data reveal a spectrum in the extent to which *A. thaliana* mitochondria contain aaRSs that are shared with the chloroplasts, aaRSs that are shared with the cytosol, or a mixture of both. Our findings indicate that ThrRS, ValRS, and (to a lesser extent) AlaRS are closer to the end of the spectrum of sharing between mitochondria and the cytosol than previously appreciated (Figures 2-3, Table 2).

### Changes in aaRS targeting and tRNA interactions associated with massive mitochondrial tRNA gene loss in Silene conica

With only two tRNA genes (tRNA-Ile and tRNA-fMet), the *S. conica* mitogenome represents one of the most extreme cases of mitochondrial tRNA gene loss in plants (Sloan, Alverson, Chuckalovcak, et al. 2012), with potential widespread effects on aaRS-tRNA interactions. The functional replacement of mitochondrial tRNA genes via import of their cytosolic counterparts raises the possibility that *S. conica* mitochondria also import the corresponding cytosolic aaRSs rather than rely on the ancestral organellar aaRSs (Warren et al. 2023). To assess whether *A. thaliana* and *S. conica* differ in their usage of organellar-like vs cytosolic-like aaRSs in their mitochondria, we devised a metric that represents the balances between these aaRS types on a 0 (all organellar) to 1 (all cytosolic) scale based on PSM counts in the mitochondrial fraction (see Figure 4 and Methods, as well as Figure S7 for an equivalent analysis based on ion intensities).

**Figure 4.**
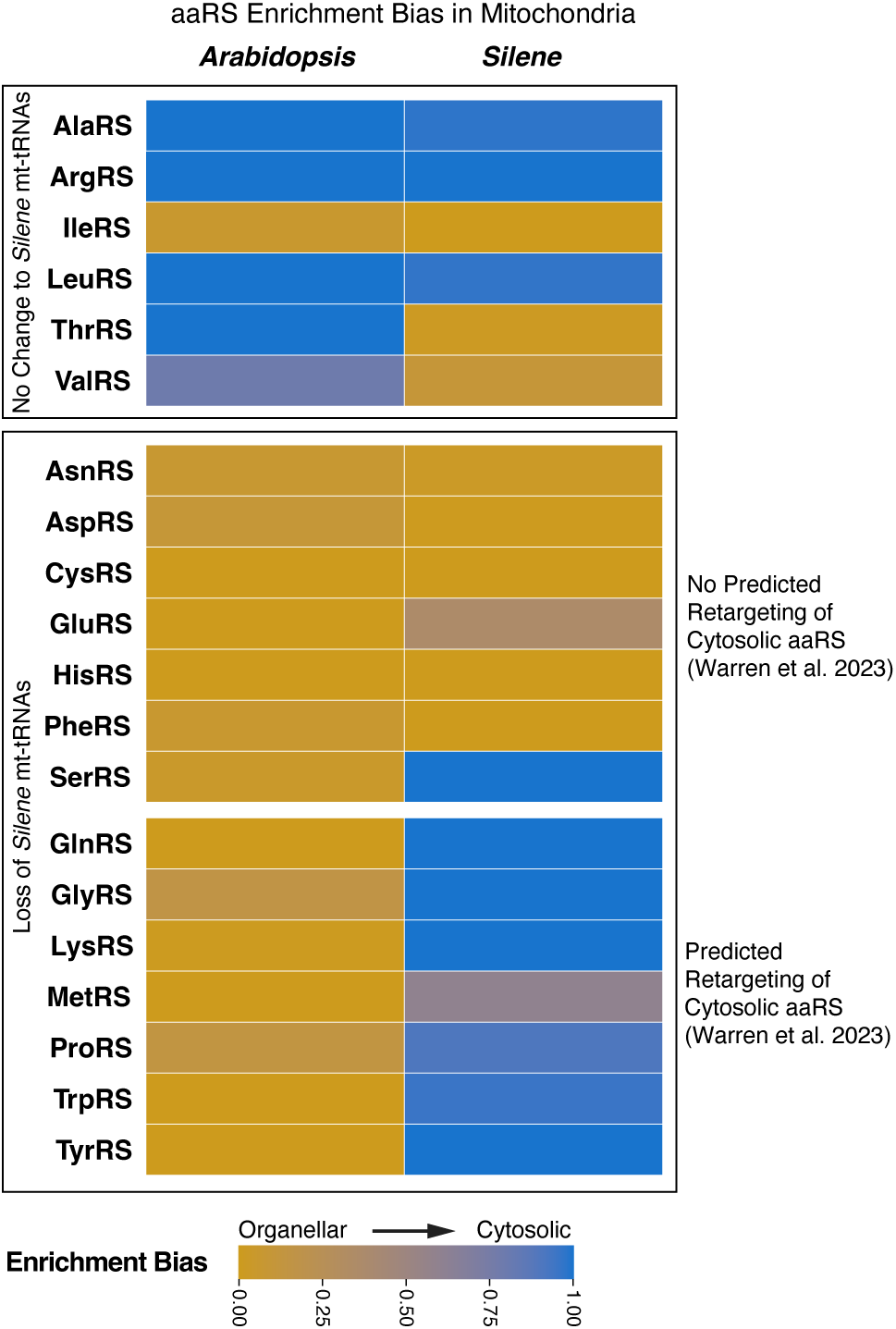
Summary of whether mitochondrial samples were biased towards containing organellar-like vs. cytosolic-like aaRSs based on proteomic analysis. The mitochondrial enrichment bias metric was calculated as described in the Methods. Note that there is no organellar-like GlnRS because mitochondria and plastids use GluRS and the indirect aminoacylation pathway for tRNA-Gln (Pujol et al. 2008). Therefore, enrichment bias for GlnRS was simply reported as “organellar” if the cytosolic-like GlnRS was not detected in mitochondria and cytosolic if the cytosolic-like GlnRS was detected in mitochondria. Overall, the results strongly align with previous predictions (Warren et al. 2023) for which cytosolic aaRSs were or were not retargeted to mitochondria with the exception of SerRS, which showed clear evidence for retargeting despite the lack of any predicted transit peptide. A similar analysis based on ion intensity is available in Figure S7.

Only six of the 20 aaRS families are not expected to be affected by recent mitochondrial tRNA gene loss in the *Silene* lineage – either because *S. conica* has retained the corresponding tRNA gene in its mitochondrial genome (IleRS) or because the corresponding tRNA gene had already been lost prior to the most recent common ancestor of angiosperms (AlaRS, ArgRS, LeuRS, ThrRS, and ValRS). Note that even though the *S. conica* mitogenome retains a gene for the initiator tRNA-fMet, it has lost the elongator tRNA-Met gene, so interactions involving MetRS may still have been perturbed. For four of these six aaRS families (AlaRS, ArgRS, IleRS, and LeuRS), mitochondria from *A. thaliana* and *S. conica* exhibit the same bias with respect to cytosolic-like vs. organellar-like aaRSs based on the aforementioned metric (Figure 4). However, the bias towards cytosolic-like ThrRS and ValRS in *A. thaliana* mitochondria (see preceding section) is not shared by *S. conica*. In these cases, it is likely that *S. conica* has retained the ancestral state (an organellar-like aaRS in the mitochondria) and that the *Arabidopsis* lineage has evolved to import a cytosolic-like enzyme.

The other 14 aaRS types are potentially affected by the large-scale loss/replacement of mitochondrial tRNA genes in *S. conica*. Previous analysis based on *in silico* prediction and fusions of putative transit peptides to GFP (Warren et al. 2023) found that a cytosolic-like aaRS likely gained targeting to the mitochondria in seven of these 14 cases (GlnRS, GlyRS, LysRS, MetRS, ProRS, TrpRS, and TyrRS) where a native mitochondrial tRNA gene was replaced by import of its cytosolic counterpart. Proteomic data supported these predictions in all seven cases (Figure 4), indicating that the ancestral pairing of cytosolic-like tRNAs and aaRSs has likely been preserved and simply retargeted to also function in the mitochondria. Notably, cytosolic-like and organellar-like MetRS enzymes were both detected in *S. conica* mitochondria (Figure 4), which might reflect the contrasting pattern of loss/replacement of the elongator tRNA-Met gene but retention of the initiator tRNA-fMet gene in the mitogenome (Sloan, Alverson, Chuckalovcak, et al. 2012). As such, we would speculate that the organellar-like MetRS charges the tRNA-fMet that is still encoded in the *S. conica* mitogenome, whereas the cytosolic-like MetRS charges a tRNA-Met that is imported from the cytosol to functionally replace the mitochondrially encoded elongator tRNA-Met gene.

The remaining seven aaRS families (AsnRS, AspRS, CysRS, GluRS, HisRS, PheRS, and SerRS) were not previously predicted to show retargeting of the cytosolic-like aaRS in *S. conica* despite importing the cytosolic tRNA counterparts (Warren et al. 2023). Our analysis here generally supported these aaRS targeting predictions (Figure 4), implying that these organellar-like aaRSs are now responsible for charging the imported cytosolic-like tRNAs. However, SerRS was a notable exception, showing clear proteomic evidence of retargeting of the cytosolic-like SerRS to the mitochondria (Figure 4). This finding resolves a mystery because previous transit peptide analysis had suggested that the ancestral organellar-like SerRS had lost targeting to mitochondria and was localized exclusively to chloroplasts in the *Silene* lineage, making it unclear which enzyme was providing SerRS function in mitochondria (Warren et al. 2023). Our proteomic results indicate that SerRS represents an eighth case of retargeting a cytosolic-like aaRS to the mitochondria. Our analysis also detected a mix of both organellar-like and cytosolic-like GluRS proteins in *S. conica* mitochondria (Figure 4). It is possible that the cytosolic-like protein is a truncated version of the enzyme that was previously predicted based on full-length mRNA sequencing and *in silico* targeting analysis (Warren et al. 2023). If so, it is not clear whether this protein would have a functional role in aminoacylation in the mitochondria given that it lacks a substantial N-terminal portion (129 amino acids) of the enzyme body. For the remaining five of these aaRSs (AsnRS, AspRS, CysRS, HisRS, and PheRS), it is likely that ancestral organellar enzyme remains primarily or solely responsible for aminoacylation in the mitochondria, although we cannot fully rule out a role of their cytosolic counterparts despite little or no detection of them in the *S. conica* mitochondrial fractions.

### Duplication and subfunctionalization of the organellar PheRS in Silene conica

PheRS is one of the aaRS types that shows no evidence of cytosolic enzyme retargeting despite the loss of the mitochondrial tRNA-Phe and apparent functional replacement with its cytosolic counterpart (Figure 4) (Warren et al. 2023). The persistence of the organellar-like PheRS in *S. conica* mitochondria follows a broader evolutionary pattern, as this aaRS appears to be one of the very last to be functionally replaced in the mitochondria even in cases of complete loss of all tRNA genes from the mitogenome (Pett and Lavrov 2015; DeTar et al. 2024). The recalcitrance of the organellar-like PheRS is thought to result from its cytosolic counterpart being divided into two subunits, making it improbable for both subunits to independently gain import into the mitochondria and serve as a viable functional replacement. In the *Silene* lineage, the organellar-like PheRS gene has been duplicated, and fusion of putative transit peptides to GFP suggested that the paralogs have subfunctionalized with one specializing on the mitochondria and the other specializing on the chloroplasts (Warren et al. 2023). Our proteomic analysis supported this subfunctionalization model, as the *S. conica* chloroplast fraction only yielded PSMs for the putative chloroplast specialist, whereas the mitochondrial fraction was dominated by PSMs from the putative mitochondrial specialist (Figure 5A). Ion intensities mirrored this pattern and showed a significant difference in the relative abundance of the two duplicates between the mitochondrial and chloroplast fractions (*p* = 0.0029; Table S2).

**Figure 5.**
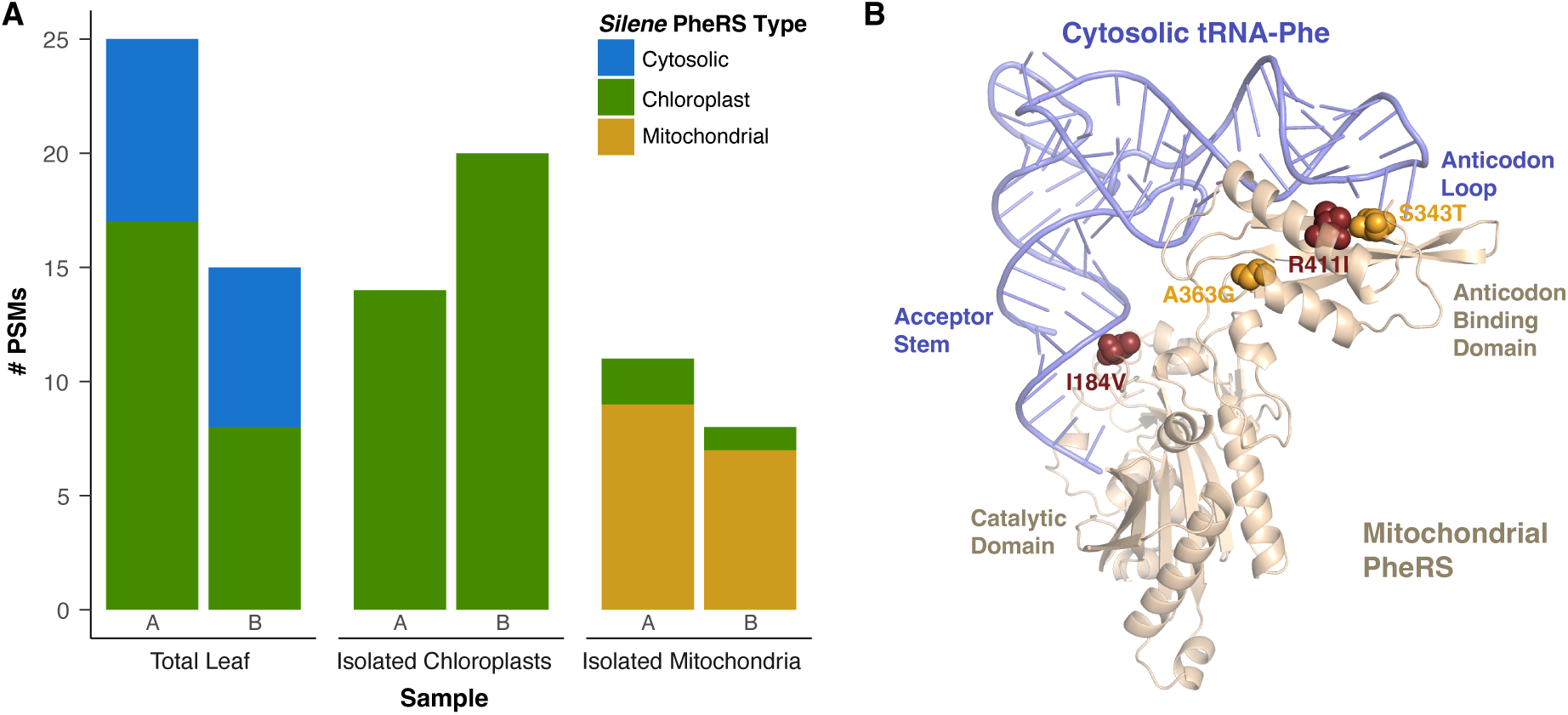
Duplication of organellar PheRS has led to subfunctionalized copies with specialized chloroplast and mitochondrial localization. (A) PSM counts in *Silene conica* samples (with A and B replicates shown separately for each sample type) indicate that the putative chloroplast PheRS was the only one present in the chloroplast fraction, whereas the putative mitochondrial type dominated the mitochondrial fraction, aligning with previous predictions (Warren et al. 2023). (B) AlphaFold3 model of the structural interaction between *S. conica* mitochondrial PheRS and cytosolic tRNA-Phe. Four substitutions at PheRS residues that are otherwise broadly conserved across green plants are labeled and highlighted with “sphere” representation. Two of those substitutions (I184V) and (R411I) are emphasized with a darker color because the exact same substitutions evolved in parallel in another angiosperm (*Sapria himalayana*) with an organellar-like PheRS that is expected to function exclusively in the mitochondria and charge imported cytosolic tRNA-Phe.

The organellar (mitochondrial) PheRS in humans has been found to have the capacity to charge tRNA-Phe substrates from diverse organisms (Klipcan et al. 2012). If plant organellar PheRSs share this capacity, it may have predisposed them to charge imported cytosolic tRNA-Phe in the mitochondria of *S. conica*. In addition, the division of labor observed between the two organellar-like PheRSs in *S. conica* could be a response to the challenges associated with a single organellar-like aaRS having to recognize both a cyanobacterial-like tRNA-Phe in chloroplasts and a cytosolic-like tRNA-Phe imported into the mitochondria.

To identify amino acid substitutions that might potentially have improved the ability of the *S. conica* mitochondrial PheRS to charge cytosolic tRNA-Phe, we compared the *S. conica* organellar PheRS sequences to each other and to orthologs from a broad sampling of species that represented angiosperms (including eudicots, monocots, magnoliids, and *Amborella*), gymnosperms (*Ginkgo*), bryophytes (*Physcomitrium*), and green algae (*Micromonas* and *Ostreococcus*). The enzyme bodies (i.e., after removal of putative transit peptide sequences) of the *S. conica* mitochondrial and chloroplast PheRSs differed by 39 amino acid substitutions (out of 374 positions). Four of the substitutions in the mitochondrial PheRS (I184V, S343T, A363G, and R411I) were at sites that were otherwise universally conserved across our sampling of plant and algal taxa (Figure S8A). No substitutions were found at any such conserved sites in the *S. conica* chloroplast PheRS sequence.

To further explore the potential relevance of these four substitutions, we compared the PheRS sequence to a representative of the Rafflesiaceae (*Sapria himalayana*) because this family of parasitic (non-photosynthetic) plants has also lost the native tRNA-Phe gene from its mitogenome, and it has lost its plastid genome entirely (Molina et al. 2014; Smith and Asmail 2014; DeTar et al. 2024). Therefore, its organellar PheRS is expected to function exclusively in the mitochondria (due to lack of translation in the plastid) and likely charges imported cytosolic tRNA-Phe. Strikingly, the *S. himalayana* PheRS has independently evolved the exact same amino acid substitutions at two of the four sites identified in the *S. conica* mitochondrial PheRS (I184V and R411I; Figure S8A). The R411I substitution is at a residue predicted to interact with the G37 position in cytosolic tRNA-Phe (Figure 5B), which is situated immediately adjacent to the anticodon and represents the only sequence difference in the anticodon loop between cytosolic and mitochondrial tRNA-Phe (Figure S8B). The I184V substitution is at a position abutting the 3′ side of the tRNA-Phe acceptor stem (Figure 5B), which contains multiple nucleotide substitutions that distinguish cytosolic and mitochondrial tRNA-Phe (Figure S8B). We also compared to the organellar PheRS from another parasitic plant (*Balanophora fungosa*) that has lost the mitochondrial tRNA-Phe gene. The *B. fungosa* organellar PheRS is expected to function in both the mitochondria and plastids because the Balanophoraceae lineage still retains a plastid genome (Su et al. 2019; Ceriotti et al. 2021). However, this plastid genome has also lost its copy of the tRNA-Phe gene, so the *B. fungosa* organellar PheRS is expected to exclusively charge cytosolic tRNA-Phe. We found that *B. fungosa* PheRS also carried the R411I substitution. It also had a substitution at residue 184, although it was an Ile-to-Thr change rather than the Ile-to-Val substitution observed in *S. conica* and *S. himalayana*.

Given the recurrence of substitutions in *S. conica*, *S. himalayana*, and *B. fungosa* at these two sites that are otherwise highly conserved in green plants, we speculate that they are involved in the specialization of these enzymes to charging an imported cytosolic tRNA-Phe substrate. In the *S. conica* lineage, it appears that these substitutions occurred prior to the divergence of multiple *Silene* species that were previously sampled (Warren et al. 2023). However, another member of this family (*Agrostemma githago*) shares the organellar PheRS paralogs with *Silene* (Warren et al. 2023), but it does not share these two amino acid substitutions. Because *A. githago* independently lost the mitochondrial tRNA-Phe gene and must use its imported cytosolic counterpart (Warren et al. 2021), the lack of these two substitutions in the *A. githago* mitochondrial PheRS implies that they are not essential for charging cytosolic tRNA-Phe. Likewise, it is possible that other amino acid substitutions distinguishing the *S. conica* mitochondrial and plastid PheRSs might contribute to their specialization on different tRNA substrates even if they are not at positions that are widely conserved in other taxa. Furthermore, it is possible this gene duplication has occurred for reasons that are entirely unrelated to changes in tRNA substrates. Performing aminoacylation assays with recombinantly expressed proteins (Gamper and Hou 2020) would be a promising route to assess the effect of specific substitutions on charging efficiency with different tRNA substrates and to distinguish among these alternative hypotheses.

### Loss of the GatCAB complex in Silene conica mitochondria

Like most bacteria, plant mitochondria and plastids generally lack GlnRS activity and instead use an indirect pathway in which GluRS indiscriminately charges tRNA-Gln with Glu followed by enzymatic conversion of Glu to Gln by the glutamyl-tRNA amidotransferase enzyme complex (GatCAB) (Pujol et al. 2008). In *S. conica*, the loss of the mitochondrial tRNA-Gln gene has been associated with import of cytosolic-like tRNA-Gln and GlnRS into the mitochondria (Figure 2). Therefore, we would predict that GatCAB activity is no longer necessary in *S. conica* mitochondria because they now use the direct Gln aminoacylation pathway typical of cytosolic translation (Rogers and Söll 1995). Our proteomic analysis supported this prediction. All three GatCAB subunits were detected in both the mitochondria and chloroplast fractions from *A. thaliana* but only in the chloroplast fraction from *S. conica* (Table S3). Therefore, the GatCAB complex and the indirect Gln aminoacylation pathway appears to no longer function in *S. conica* mitochondria. However, it is possible that the GatCAB complex is still targeted to *S. conica* mitochondria at low levels that were difficult to detect in our analysis but are nonetheless biologically relevant.

### Retention of other components of tRNA metabolism machinery in Silene conica mitochondria

The two remaining tRNA genes in the *S. conica* mitogenome (tRNA-fMet and tRNA-Ile) are noteworthy because they both have distinctive bacterial-like features. Bacterial translation is initiated with an N-formylmethionine (fMet), which is synthesized by the methionyl-tRNA formyltransferase (MTF). After tRNA-fMet is initially charged with Met, the MTF enzyme formylates the amino group of the Met residue to produce fMet (Ibba and Söll 2004). Meanwhile, a class of bacterial tRNA-Ile genes have a CAT anticodon, which would typically correspond to an ATG (Met) codon. However, the C base in this anticodon is modified to lysidine by tRNA-Ile lysidine synthetase (TilS), which results in decoding of Ile codons (Suzuki and Miyauchi 2010). Homologs of bacterial MTF and TilS are both found in plant nuclear genomes and expected to function in mitochondrial and plastid translation systems (Warren and Sloan 2020). Given the retention of tRNA-fMet and tRNA-Ile genes in the *S. conica* mitogenome, we would predict that MTF and TilS are also functional in *S. conica* mitochondria. Accordingly, we detected both proteins in the *S. conica* mitochondrial fraction (Table S3). TilS was only supported by a single PSM in one of the *S. conica* mitochondrial samples, which is likely due to low overall expression, as we did not detect it in chloroplast or total-leaf samples from *S. conica* or in any *A. thaliana* samples. A previous deep proteomic analysis of *A. thaliana* mitochondria did detect TilS, albeit at very low abundance (Rugen et al. 2024). Therefore, the detection of both these enzymes in *S. conica* mitochondria suggests that the bacterial-like translation features associated with tRNA-fMet and tRNA-Ile have been retained, which contrasts with the apparent functional loss of GatCAB and many aaRSs.

Mitochondrial tRNA metabolism also relies on additional enzymes that are typically shared with other subcellular compartments (von Braun et al. 2007; Canino et al. 2009; Gobert et al. 2010; Gutmann et al. 2012; Salinas-Giegé et al. 2015), including those responsible for cleaving primary transcripts to remove 5′ ends (protein-only RNase P [PRORP]) or 3′ ends (tRNase Z) and for adding a 3′ CCA tail (CCAse). In general, we had limited sensitivity to detect these enzymes with our dataset (Table S3). Although we did confirm the expected presence of CCAse in *S. conica* mitochondria, the general lack of signal for PRORP or tRNase Z enzymes across all samples precludes any interpretation of whether there has been retargeting of these enzymes associated with loss of mitochondrial tRNA genes.

### Mitochondrial ribosomal subunit gene loss, transfer, and replacement in Silene conica

The *S. conica* mitogenome has lost most of the genes that encode ribosomal protein subunits, retaining only three (*rpl5*, *rps3*, and *rps13*) of the 15 that were present in most recent common ancestor of angiosperms (Adams, Qiu, et al. 2002; Kubo and Arimura 2010; Sloan, Alverson, Chuckalovcak, et al. 2012). Although *rps3* was originally annotated as a pseudogene in the *S. conica* mitogenome due to the apparent loss of the first exon and the presence of large indels (Sloan, Alverson, Chuckalovcak, et al. 2012), we detected Rps3 peptides via LC-MS/MS, indicating that it remains an expressed, functional gene. More generally, our proteomic dataset provides potential insights into how the extensive gene loss from the mitogenome occurred without disrupting function of the mitochondrial ribosome (mitoribosome). Our results point to diverse mechanisms by which these genes were replaced (Figure 6; Table 3).

**Figure 6.**
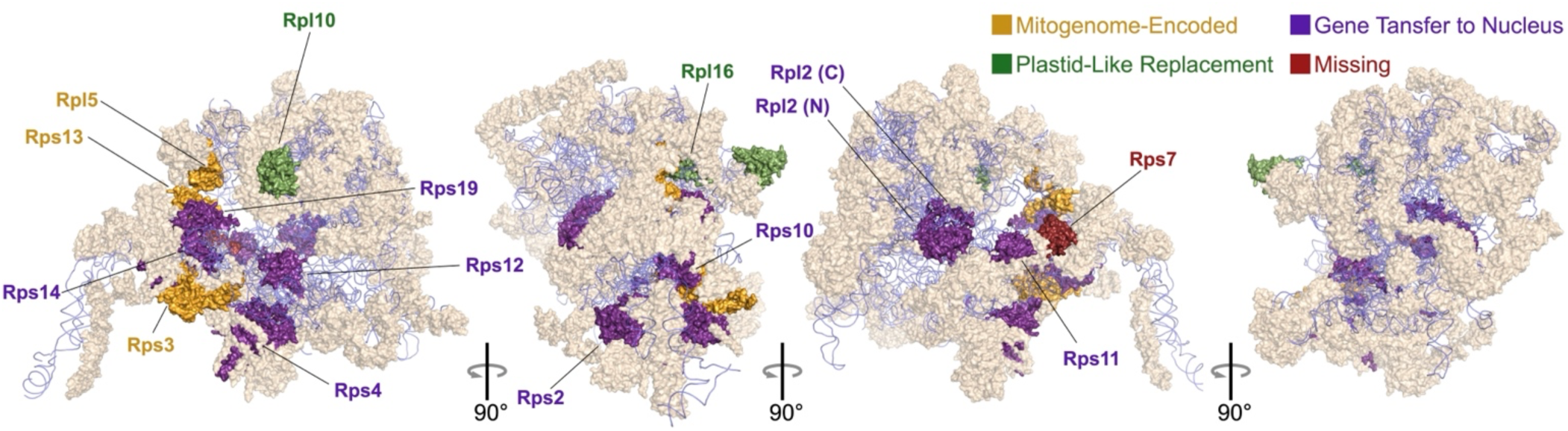
Structure of *A. thaliana* mitoribosome (PDB accession 6XYW; Waltz et al. 2020). Ribosomal proteins are shown with surface renderings, and rRNAs are shown in blue with cartoon renderings. Subunits that were encoded by genes in the ancestral angiosperm mitogenome are highlighted and colored according to their evolutionary history in *S. conica*. Mitochondrial localization of these subunits in *S. conica* is supported by our LC-MS/MS data (Tables 3 and S4). However, we have not directly tested for assembly of these *S. conica* proteins into the mitoribosome itself. Note that the Rps1 subunit is not pictured because it has been lost in *A. thaliana* (Skaltsogiannis et al. 2024) even though the gene is retained in most angiosperms, including the copy that has been transferred to the nucleus in *S. conica* (Table 3).

**Table 3.**
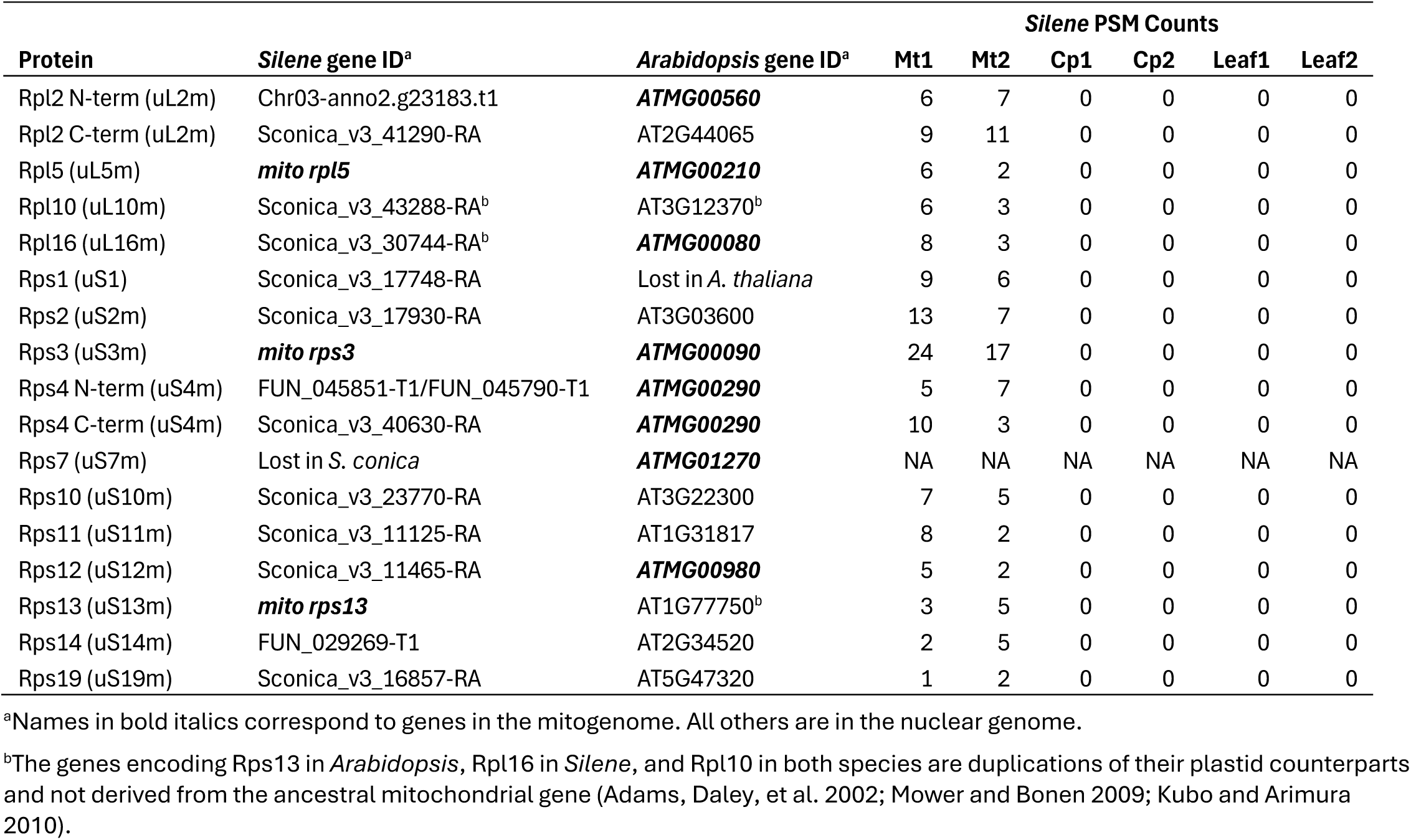
Genes encoding subunits of the mitoribosome that were ancestrally present in the angiosperm mitogenome.

| Protein | *Silene* gene ID<sup>a</sup> | *Arabidopsis* gene ID<sup>a</sup> | *Silene* PSM Counts |  |  |  |  |  |
| --- | --- | --- | --- | --- | --- | --- | --- | --- |
|  |  |  | Mt1 | Mt2 | Cp1 | Cp2 | Leaf1 | Leaf2 |
| Rpl2 N-term (uL2m) | Chr03-anno2.g23183.t1 | **ATMG00560** | 6 | 7 | 0 | 0 | 0 | 0 |
| Rpl2 C-term (uL2m) | Sconica_v3_41290-RA | AT2G44065 | 9 | 11 | 0 | 0 | 0 | 0 |
| Rpl5 (uL5m) | **mito rpl5** | **ATMG00210** | 6 | 2 | 0 | 0 | 0 | 0 |
| Rpl10 (uL10m) | Sconica_v3_43288-RA<sup>b</sup> | AT3G12370<sup>b</sup> | 6 | 3 | 0 | 0 | 0 | 0 |
| Rpl16 (uL16m) | Sconica_v3_30744-RA<sup>b</sup> | **ATMG00080** | 8 | 3 | 0 | 0 | 0 | 0 |
| Rps1 (uS1) | Sconica_v3_17748-RA | Lost in *A. thaliana* | 9 | 6 | 0 | 0 | 0 | 0 |
| Rps2 (uS2m) | Sconica_v3_17930-RA | AT3G03600 | 13 | 7 | 0 | 0 | 0 | 0 |
| Rps3 (uS3m) | **mito rps3** | **ATMG00090** | 24 | 17 | 0 | 0 | 0 | 0 |
| Rps4 N-term (uS4m) | FUN_045851-T1/FUN_045790-T1 | **ATMG00290** | 5 | 7 | 0 | 0 | 0 | 0 |
| Rps4 C-term (uS4m) | Sconica_v3_40630-RA | **ATMG00290** | 10 | 3 | 0 | 0 | 0 | 0 |
| Rps7 (uS7m) | Lost in *S. conica* | **ATMG01270** | NA | NA | NA | NA | NA | NA |
| Rps10 (uS10m) | Sconica_v3_23770-RA | AT3G22300 | 7 | 5 | 0 | 0 | 0 | 0 |
| Rps11 (uS11m) | Sconica_v3_11125-RA | AT1G31817 | 8 | 2 | 0 | 0 | 0 | 0 |
| Rps12 (uS12m) | Sconica_v3_11465-RA | **ATMG00980** | 5 | 2 | 0 | 0 | 0 | 0 |
| Rps13 (uS13m) | **mito rps13** | AT1G77750<sup>b</sup> | 3 | 5 | 0 | 0 | 0 | 0 |
| Rps14 (uS14m) | FUN_029269-T1 | AT2G34520 | 2 | 5 | 0 | 0 | 0 | 0 |
| Rps19 (uS19m) | Sconica_v3_16857-RA | AT5G47320 | 1 | 2 | 0 | 0 | 0 | 0 |

<sup>a</sup>Names in bold italics correspond to genes in the mitogenome. All others are in the nuclear genome.
<sup>b</sup>The genes encoding Rps13 in *Arabidopsis*, Rpl16 in *Silene*, and Rpl10 in both species are duplications of their plastid counterparts and not derived from the ancestral mitochondrial gene (Adams, Daley, et al. 2002; Mower and Bonen 2009; Kubo and Arimura 2010).

Nine of the twelve genes that have been lost (*rpl2*, *rps1*, *rps2*, *rps4*, *rps10*, *rps11*, *rps12*, *rps14*, and *rps19*) have likely followed the typical route of intracellular gene transfer from the mitogenome to the nucleus (Adams and Palmer 2003). In two of these cases (*rpl2* and *rps4*), the transfer appears to have occurred in two pieces, resulting in different nuclear genes encoding separate, non-overlapping portions of each subunit (Table 3). In the case of *rpl2*, the splitting of the gene and transfer of the portion encoding the C-terminal end of the protein were ancient events that appear to have occurred in a common ancestor of the core eudicots (Adams et al. 2001). Whereas the gene encoding the N-terminal portion of the protein is retained in the *A. thaliana* mitogenome, it too has been transferred to the nucleus in the *S. conica* lineage (Table 3), in addition to independent transfers in multiple other eudicot lineages (Adams et al. 2001).

Two more genes absent from the *S. conica* mitogenome (*rpl10* and *rpl16*) also appear to have been functionally replaced through import of a nuclear-encoded protein. However, in these cases, the nuclear gene is plastid-like rather than an intracellular transfer of the mitochondrial gene itself. The nuclear gene encoding the plastid-targeted Rpl10 subunit in *S. conica* has been duplicated, with one copy now targeted exclusively to the mitochondria based on our LC-MS/MS data. Although many angiosperms retain the *rpl10* gene in the mitogenome, similar duplications and neofunctionalization of plastid-targeted homologs were previously identified in monocots and in the Brassicaceae (Mower and Bonen 2009; Kubo and Arimura 2010). In the case of *rpl16*, the *S. conica* nuclear genome contains a plastid-like gene copy encoding a protein that we exclusively detected in the mitochondrial fraction, and the *S. conica* plastid genome still contains a typical *rpl16* gene. As such, it appears that a copy of the plastid *rpl16* gene was transferred to the nucleus but gained targeting to the mitochondria, facilitating the loss of its mitochondrial homolog. This scenario differs from the case of *rpl10* because there is no indication that the transferred *rpl16* gene was ever targeted back to the plastid. Therefore, this evolutionary transition may have required overcoming multiple barriers almost simultaneously. Specifically, the transferred nuclear gene would have had to gain expression and targeting to the mitochondria but also adapt to function in the novel context of the mitoribosome.

The one other ribosomal gene that has been lost from the *S. conica* mitogenome (*rps7*) has no detectable homolog in the nuclear genome other than the distantly related family member that encodes a subunit of the cytosolic ribosome (*RPS5*; Sconica_v3_48856-RA). The plastid *rps7* homolog is still retained in the plastid genome itself. Across angiosperm diversity, the *rps7* gene has been subject to an unusually large number of losses from the mitogenome (Adams, Qiu, et al. 2002). Although there are cases where loss of the mitochondrial copy of *rps7* appears to have been accompanied by transfer to the nucleus (Liu et al. 2009), outright loss of the gene may also be common (Adams, Qiu, et al. 2002). The *S. conica* cytosolic-like homolog (Sconica_v3_48856-RA) was detectable in all three sample types. However, proteomic detection in the mitochondrial fraction should not be taken as strong evidence for function within the mitochondria given the extremely high abundance of cytosolic ribosomes and the propensity for these ribosomes to adhere to outer mitochondrial membranes (Chang et al. 2024; Dimnet et al. 2024). Therefore, it is not clear if or how the Rps7 subunit has been replaced in the mitoribosome.

Even in angiosperms that have retained a larger number of ribosomal genes in their mitogenomes than *S. conica*, the majority of mitoribosome subunits are encoded by nuclear genes. The composition of the *A. thaliana* mitoribosome has been thoroughly characterized (Rugen et al. 2019; Waltz et al. 2019; Waltz et al. 2020; Skaltsogiannis et al. 2024), so we used proteomic data from the *S. conica* mitochondrial fraction to infer whether these subunits are conserved in the *S. conica* mitoribosome. Of the 70 nuclear-encoded subunits of the *A. thaliana* mitoribosome (Table S4), only bL32m (AT1G26740) lacks a detectable ortholog among the annotated *S. conica* protein set, and this appears to be an annotation issue because homologous sequence is detectable with a TBLASTN search against the *S. conica* nuclear genome. Only three of the annotated proteins (bS21m, mS38, and bTHXm) were not detected in any of our samples, and every one of the proteins that was detected had higher PSM counts in the mitochondrial fraction than in chloroplast or total leaf samples. Indeed, more than 90% of these proteins were exclusively found in the mitochondrial fraction based on PSM counts, and even with only two biological replicates, the majority of these putative mitoribosome subunits showed significant enrichment in the *S. conica* mitochondrial fraction relative to total leaf tissue based on normalized ion intensities (Table S4). Therefore, orthologous gene content and mitochondrial proteomes suggest a high degree of stability in the ancestral nuclear-encoded components of the *S. conica* mitoribosome. However, confirming that these ancestral components, as well as subunits resulting from inferred gene transfers and replacements, are actually assembled as part of the ribosome complex will require purification and structural analysis of the mitoribosomes themselves.

### Limitations of this study

This study offers the first direct characterization of the *S. conica* mitochondrial and chloroplast proteomes, which had only previously been inferred from genomic/transcriptomic analysis, *in silico* predictions, and use of fluorescent reporters fused to putative transit peptides. However, the study has multiple limitations that can be addressed with future efforts to provide a more comprehensive characterization of these proteomes. First, we were only able to incorporate two biological replicates into the design, limiting statistical power for tests of enrichment and differential proteome composition. Second, tissue collection and sample preparations were spread out across multiple days (see Methods) to facilitate large scale organelle purification steps, but it would be preferable to obtain more precise pairing of sample types for comparative purposes. Third, we only used trypsin digests, whereas parallel analyses with multiple proteases has been found to increase representation of plant mitochondrial proteomes (Rugen et al. 2024). Fourth, numerous dimensions of proteome variation remain unexplored in this study, including variation among tissues, cell types, environmental conditions, developmental stages, organelle types (e.g., proplastids vs mature chloroplasts), and sub-organellar fractions (e.g., nucleoids, ribosomes, thylakoids, etc) – all of which represent interesting areas for future investigation.

## Conclusions

The ongoing loss of angiosperm mitochondrial genes has been well characterized for more than two decades, with especially pronounced effects on the genes encoding translational machinery such as tRNAs and ribosomal proteins (Adams, Qiu, et al. 2002; Richardson et al. 2013). However, direct proteomic analysis of mitochondrial translation machinery has been lacking in species such as *S. conica* that show extreme reductions in mitochondrial gene content. Our study details the extensive changes that accompany mitochondrial gene loss, highlighting alternative evolutionary pathways for functionally replacing genes and responding to perturbations in the network of tRNA-interacting enzymes.

For example, our work confirms that numerous cytosolic-like aaRSs have been retargeted to the mitochondria in *S. conica* (Figure 4), presumably preserving ancestral charging relationships with cytosolic tRNAs that are newly imported into the mitochondria (Warren et al. 2023). We found that this set of retargeted cytosolic-like aaRSs includes SerRS even though *in silico* predictions suggested that it has not gained an N-terminal extension that could serve as a mitochondrially targeting transit peptide (Warren et al. 2023). This finding illustrates why direct proteomic analysis is an important complement to *in silico* predictions and assays based on fusing reporters to N-terminal peptides (Mireau et al. 1996; Uwer et al. 1998; Souciet et al. 1999; Peeters et al. 2000; Duchêne et al. 2001; Duchêne et al. 2005; Warren et al. 2023). On the other hand, our work also shows that many cytosolic-like aaRSs were likely not retargeted to the mitochondria despite import of their cognate cytosolic tRNAs. Therefore, organellar-like aaRSs are presumably charging novel substrates (cytosolic-like tRNAs) in these cases.

Likewise, functional replacement of mitochondrial ribosomal protein genes appears to have followed multiple pathways (Figure 6), including relocation of the gene to the nucleus, replacement by a plastid-like counterpart, or (in the case of *rps7*) outright loss or replacement with a subunit lacking detectable homology. Overall, the recent evolutionary changes in *S. conica* highlight that the composition of plant mitochondrial translation machinery is still highly dynamic despite billions of years since the establishment of mitochondria in the eukaryotic lineage. Although the processes of mitochondrial gene loss, transfer, and replacement have been unusually extensive and recent in the *S. conica* lineage, they are all common themes in angiosperms (Adams, Qiu, et al. 2002; Warren and Sloan 2020) that have also contributed to variation in mitochondrial translation machinery more broadly across the tree of life.

## Materials and Methods

### Plant growth and organelle isolations

*Arabidopsis thaliana* Col-0 seeds were stratified at 4 °C in water for 3 days prior to sowing in 3-inch pots with 5 seeds per pot. *Agrostemma githago* KEW0053084 (Warren et al. 2023) and *S. conica* ABR (Fields et al. 2023) were sown in 4-inch pots with 6 seeds per pot. Pots contained Pro-Mix BX potting media and were covered with clear plastic domes for ∼1 week until seedlings emerged. All plants were grown on shelves with fluorescent lighting (∼100 µE m^-2^ sec^-1^) at ∼22-23 °C under short-day conditions (10-hr light / 14-hr dark). Mitochondrial and chloroplast isolations were performed with tissue harvested either ∼9 weeks after sowing (*A. thaliana* and *A. githago*) or ∼12 to 13 weeks after sowing (*S. conica*). Tissue samples for total leaf protein extraction from each species were harvested shortly before (within 1 week) the respective organelle isolations.

Two biological replicates were performed for each species. Mitochondrial and chloroplast fractions were isolated with Percoll gradients. Mitochondrial isolations were performed as described previously (Warren et al. 2021) based on a modified protocol from Meyer et al. (2009), using ∼70 g of rosette leaf tissue per replicate. Chloroplast isolations were performed using a modified version of published protocols (van Wijk et al. 2007; Kley et al. 2010). Briefly, ∼2 g of rosette leaf tissue was ground in a prechilled mortar with 20 ml of ice-cold chloroplast grinding and wash buffer (cpGW: 50mM HEPES pH 7.5, 5 mM EDTA, 0.3 M sorbitol, 10 mM NaHCO3, 0.5 mM DTT), filtered through miracloth and centrifuged at 1,300 rcf for 5 min at 4 °C. The resulting chloroplast pellet was resuspended in ∼2 ml cpGW, applied to a Percoll step gradient (40-80%) and centrifuged at 2,500 rcf in a swinging bucket rotor for 10 min at 4 °C. The chloroplast band at the 40-80 interface was removed and washed three times with cpGW buffer.

Chloroplasts were resuspended in 1 ml of cpGW buffer containing 1X Halt protease inhibitor (Thermo Scientific 78430), aliquoted and centrifuged at 1,000 rcf for 5 min at 4 °C. Detailed versions of the isolation protocols are available via GitHub (https://github.com/dbsloan/silene_proteomics). Whole leaf tissue and isolated mitochondrial/chloroplast pellets were flash frozen in liquid N_2_ and stored at -80 °C before shipment to the Proteome Exploration Laboratory at the California Institute of Technology for protein extraction, digestion, and LC-MS/MS analysis.

### Protein extraction and digestion

For leaf and purified chloroplast samples, 100 μl of lysis buffer from a PreOmics iST Kit was added and followed by processing with a PreOmics BeatBox Tissue Homogenizer (Preomics, Germany) for 10 min on the high setting. Samples were then centrifuged at 21,000 rcf for 2 min to remove the insoluble fraction. Protein concentration was evaluated using Pierce BCA Protein Assay Kit. Aliquots containing 100 μg of protein from each sample were digested with ProtiFi S-trap according to the manufacturer’s protocol. Briefly, the protein was reduced and alkylated with tris(2-carboxyethyl)phosphine and chloroacetamide and digested overnight with trypsin. Following elution, the resulting peptides were dried and resuspended, using 2% acetonitrile, 0.2% formic acid in water.

For purified mitochondria samples, which were smaller and contained less protein content, pellets were resuspended in 120 μl of 8 M urea with 50 mM HEPES. Samples were reduced with tris(2-carboxyethyl)phosphine (10 min, 60 °C) and chloroacetamide (15 min, room temperature). Samples were then treated with 2 μl 0.1 mg/ml LysC endopeptidase (Wako Pure Chemical) for 2 hr at 37 °C. Following sample dilution with 360 μl of 50 mM HEPES, 5 μl of 100 mM CaCl_2_ and 3 μl of the 0.1 mg/ml Trypsin (Pierce) were added for overnight digestion at 37 °C. The digested peptides were desalted using Pierce C18 Spin Columns according to the manufacturer’s protocol. The eluates were dried and resuspended using 2% acetonitrile, 0.2% formic acid in water.

### LC-MS/MS

For each sample, 500 ng was loaded onto a Thermo Scientific EASY-nLC 1200 connected to an Q Exactive HF Quadrupole-Orbitrap Hybrid Mass Spectrometer. Peptides were separated on an Aurora UHPLC Column (25 cm × 75 μm, 1.6 μm C18, AUR2-25075C18A, Ion Opticks) with a flow rate of 0.35 μl/min for a total duration of 160 min, including washing and re-equilibration. The gradient was composed of 2% Solvent B from the start, 2-6% B for 3.5 min, 6-25% B for 97 min, 25-40% B for 19.5 min, 40-98% B for 2 min, 98% B for 3 min, and 98-2% B for 2 min. The gradient was followed by 3 "see-saw" cycles (2% B for 3 min, 2-98% B for 2 min, 98% B for 3 min, and 98-2% B for 2 min) for cleaning, and the column was re-equilibrated at 2% B for 3 min. Solvent A consisted of 97.8% H_2_O, 2% acetonitrile, and 0.2% formic acid, and solvent B consisted of 19.8% H2O, 80% acetonitrile, and 0.2% formic acid. Peptides were ionized via electrospray ionization (NSI) at 2.0 kV. MS1 scans were acquired with a range of 350–1600 m/z at 60K resolution. The maximum injection time was 15 ms with an AGC target of 3 × 10^6^. MS2 scans were acquired at 30K resolution with a scan range of 200-2000 m/z. The maximum injection time was 45 ms with a minimum AGC target of 4.5 × 10^3^. The isolation window was 1.2 m/z, collision energy was 28 NCE, and loop count was set to 12. Mass spectrometer method modification and data collection were performed using Thermo Scientific Xcalibur software.

Data analysis was performed using Proteome Discoverer 3.1 (Service Pack 1), reference databases from the respective species (see below), and Sequest HT with Percolator validation. Minora Feature Detector was used to detect chromatographic peaks. Feature mapping was carried out allowing a maximum retention time shift of 10 minutes, with minimum signal-to-noise ratio of 5. Precursor ions were quantified according to feature area and normalized by using all peptides. Precursor mass tolerance was set to 10 ppm, and peptide fragment mass tolerance was set to 0.02 Da. Percolator FDRs were set at 0.01 (strict) and 0.05 (relaxed). Peptide FDRs were set at 0.01 (strict) and 0.05 (relaxed), with minimum peptide confidence set to high and a minimum peptide length of 6. High-confidence peptide matches were therefore controlled at a 0.01 FDR. Carbamidomethyl (C) was set as a static modification; oxidation (M) was set as a dynamic modification; dynamic N-Terminal modifications included: acetyl (protein N-term), met-loss (M), and met-loss+acetyl (M). The “Trypsin (Full)” protease settings were used, allowing for 2 missed cleavages. Sample-specific PSM counts were retrieved from a separate analysis in the same version of Proteome Discoverer that made use of only the SequestHT search engine and did not include met-loss dynamic modifications.

We further processed PSM and ion intensity data with custom scripts to exclude peptides that were shared between multiple protein groups. In addition, the normalized ion intensity estimates from Proteome Discoverer include contributions from peptides identified solely based on an MS1 peak that was not validated in the same sample with an MS2 spectrum. Because these estimates (flagged as “Peak Found”) have the potential to unreliably assign ion intensities to proteins, we also performed some analyses of intensities after removing these estimates with custom scripts (see text and figure legends).

Sequence for the *A. thaliana* reference protein database were obtained from the 2023-10 release of PeptideAtlas (van Wijk et al. 2021) and included the Araport11 (Cheng et al. 2017) nuclear-encoded protein sequences (longest isoform only) combined with the mitochondrial-encoded and plastid-encoded proteins curated by van Wijk et al. (2024). Loci in the large insertion of mitochondrial DNA in nuclear chromosome 2 (Stupar et al. 2001; Fields et al. 2022) that are annotated as functional protein-coding genes in the Araport11 database were removed from the reference to avoid ambiguity in mapping to the true mitochondrial-encoded proteins. For *S. conica*, annotated protein sequences were taken from the published nuclear (Fields et al. 2023), mitochondrial (Sloan, Alverson, Chuckalovcak, et al. 2012), and plastid genomes (Sloan, Alverson, Wu, et al. 2012). As with *A. thaliana*, annotated genes found in recent insertions of mitochondrial or plastid DNA into the nucleus were removed from the reference. We also added three aaRS protein sequences that were previously identified with full-length RNA-seq transcripts (Warren et al. 2023) but were not annotated in the published *S. conica* genome (Iso-Seq AspRS, HisRS, and ValRS). Mitochondrial-encoded and plastid-encoded protein sequences for *A. githago* were taken from published genomes (Sloan et al. 2014; Warren et al. 2021). Because no reference-quality annotation was available for the *A. githago* nuclear genome (although one has since been published; Mian and Leitch 2024), we used previously published full-length RNA-seq transcripts (Warren et al. 2023) for nuclear-encoded protein references. The longest open-reading frame for each transcript was identified with TransDecoder v5.7.1, and the corresponding protein-coding sequences with >99% identity were collapsed with CD-HIT v4.8.1 (Fu et al. 2012). This transcriptome reference for *A. githago* was then filtered to avoid duplicating annotated proteins from the mitochondrial and plastid reference genomes.

### Enrichment analysis of mitochondrial-encoded and plastid-encoded proteins

For each species, enrichment of mitochondrial-encoded and plastid-encoded proteins in the organellar fractions was calculated as the PSM count ratio in the corresponding organellar sample relative to total leaf tissue. PSM counts for the two biological replicates from each sample type were summed for these calculations. Proteins with fewer than 5 PSMs in total across all samples from a species were excluded. In cases where no PSMs were detected in the leaf tissue (but were detected in an organellar fraction), a minimum count of 1 was applied for the leaf sample to avoid dividing by 0. Enrichment ratios were also calculated based on ion intensity by using the reported abundance values from Proteome Discoverer. For these calculations, peptides were only included if they were assigned “High” confidence based on identification of a corresponding PSM in that sample (i.e., not “Peak Found”) and were not shared with other protein groups.

### Analysis of aaRSs and other tRNA metabolism enzymes

To investigate the subcellular localization of previously identified aaRSs (Duchêne et al. 2005; Duchêne et al. 2009; Warren et al. 2023), PSM enrichment ratios were calculated as described above for the organellar-encoded proteins. To assess how relative aaRS abundance within the mitochondria differed between species, we calculated a metric of enrichment bias between organellar-like and cytosolic-like counterparts. Specifically, the mitochondrial enrichment ratio (see above) for the cytosolic-like aaRS was divided by the sum of the mitochondrial enrichment ratios for the organellar-like and cytosolic-like aaRSs. Therefore, a value of 1 for this metric would indicate that only the cytosolic-like aaRS was found in the mitochondria, whereas a values of 0 would indicate that only the organellar-like aaRS was found. When both types were detected in the mitochondria, this metric produces an intermediate value that reflects the relative bias of organellar-like vs. cytosolic-like types. By using the enrichment ratios in these calculations, we were able to normalize to the total leaf samples and avoid directly comparing raw PSM counts between different proteins. We summed PSM counts in cases where there were multiple proteins or subunits for the same aaRS class. For some proteins, we found very low PSM counts in leaf tissue, resulting in large and highly variable enrichment ratios. Therefore, we capped enrichment ratios at a value of 3 to avoid obscuring signal of co-existing aaRS types based on these inflated values. We also repeated these analyses using ion intensity rather than PSM account as the measure of abundance.

PheRS and tRNA-Phe sequences were obtained via SHOOT (Emms and Kelly 2022), BLASTP searches of the NCBI RefSeq database (Pruitt et al. 2004), and previously curated datasets (Warren et al. 2021; Warren et al. 2023; DeTar et al. 2024). Alignments were performed with MAFFT v7.526 (Katoh and Standley 2013) under default parameters and visualized with Geneious (Kearse et al. 2012). The structure of the *S. conica* PheRS/tRNA-Phe complex was predicted using the AlphaFold3 Server (Abramson et al. 2024) under default parameters. The mitochondrial PheRS and cytosolic tRNA-Phe sequences were obtained from previous studies (Warren et al. 2021; Warren et al. 2023). The first 49 amino acids were removed from the PheRS sequence as a putative transit peptide.

The *A. thaliana* genes encoding GatCAB, MTF, TilS, PRORP, tRNase Z, and CCAse proteins were previously identified (von Braun et al. 2007; Pujol et al. 2008; Canino et al. 2009; Gutmann et al. 2012; Warren and Sloan 2020). We used reciprocal BLASTP searches against the annotated proteins sequences from the *S. conica* genome to identify orthologs and inferred presence of the enzymes in organellar fractions based on PSM counts as described above.

We further compared our results against other studies with deeper sampling of the *A. thaliana* mitochondrial proteome. Specifically, we used estimates of protein copies per mitochondrion in Dataset S1 from Fuchs et al. (2020) and the quantification from the All_Proteases_Average_(>0) column in Table S2 from Rugen et al. (2024). For Figure 3B, estimates of protein copies per mitochondrion were summed in cases of paralogs for the same enzyme, and they were averaged for the two subunits of the heteromeric cytosolic PheRS complex. See the Figure 3 legend for description of how special cases in the AlaRS, ArgRS, and GlnRS families were handled. The statistical significance of the relationship in Figure 3B was tested with the cor.test function with data that were log-transformed after adding 1 to each value to avoid log-transformation of 0 values.

### Analysis of mitoribosome subunits

The set of *A. thaliana* mitoribosome subunits was taken from the published structure (Waltz et al. 2020). We also included the uL1m (AT2G42710) and bL12m (AT3G06040) subunits, which were not captured in this structure presumably because they are located in highly mobile parts of the ribosome (Waltz et al. 2020). We used the mitochondrial-encoded Rps1 subunit from *Carica papaya* because this protein is known to have been lost entirely from *A. thaliana* (Skaltsogiannis et al. 2024). Reciprocal BLASTP searches were used to identify *S. conica* orthologs, and we inferred presence of subunits in the mitochondrial fraction based on PSM data as described above. Some ribosomal subunits were encoded by two or more closely related paralogs, and some reciprocal BLASTP searches failed due to this paralogy. In other cases, no PSMs were detected in any sample for the named subunit in the *A. thaliana* mitoribosome structure. In these cases, closely related paralogs were manually checked and substituted in where necessary. The mitoribosome structure was visualized with PyMol v3.1.3, using Protein Data Bank accession 6XYW (Waltz et al. 2020).

## Data Availability

LC-MS/MS data have been deposited to the ProteomeXchange Consortium via the PRIDE (Pérez-Riverol et al. 2024) partner repository with the dataset identifier PXD071069. Code and processed data are available via GitHub (https://github.com/dbsloan/silene_proteomics).

## Acknowledgements

We thank Baiyi Quan and the Proteome Exploration Laboratory at the California Institute of Technology for performing LC-MS/MS runs and data processing, as well as three anonymous reviewers for their highly constructive comments on a previous version of the manuscript. This work was supported by a grant from the National Science Foundation (MCB-2048407) and an HHMI Hanna H. Gray Fellowship.

---

**Figure S1.**
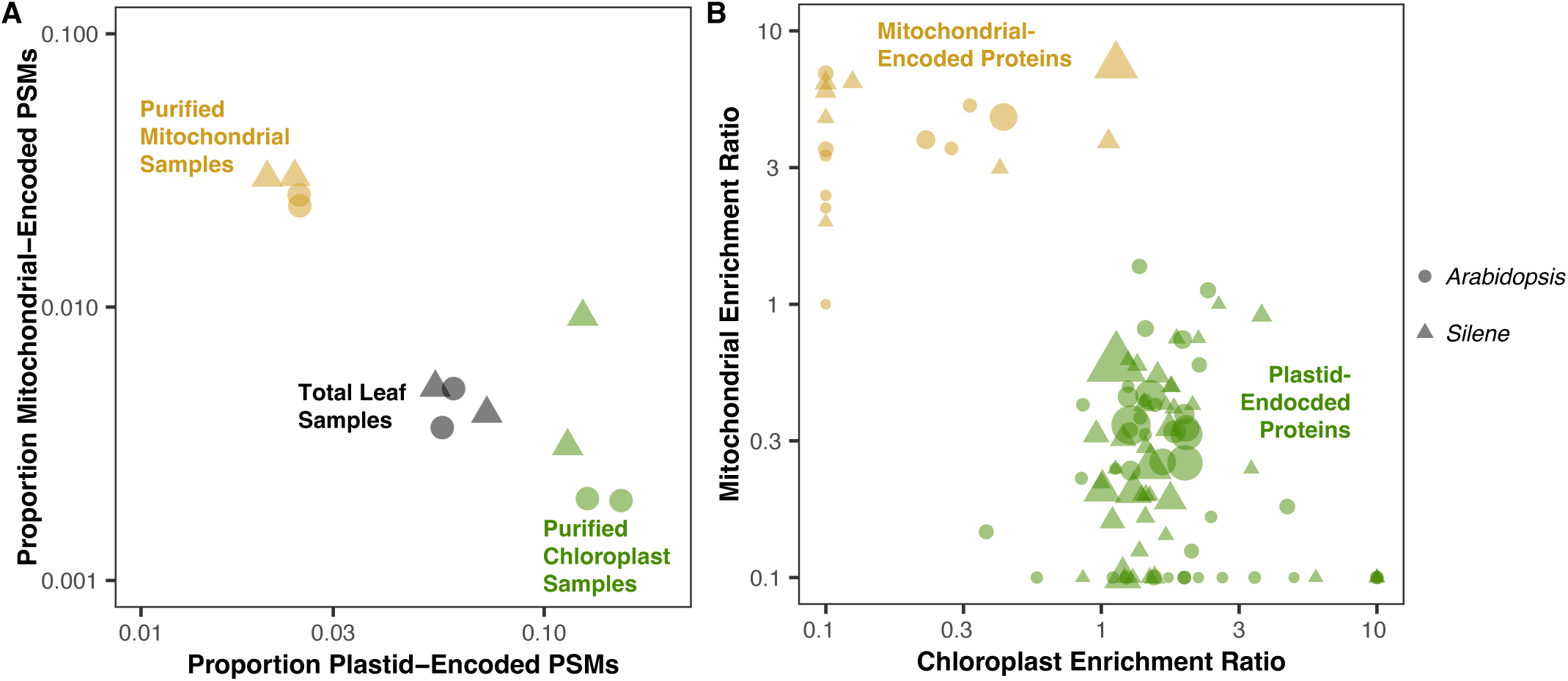
Enrichment of proteins encoded by the mitochondrial and plastid genomes in purified organelle fractions. (A) In this panel, each point represents an individual biological sample, showing its cumulative number of PSMs across all plastid-encoded proteins (x-axis) and mitochondrial-encoded proteins (y-axis) expressed as a proportion of all PSMs in the sample with each biological replicate shown separately. (B) In this panel, each point represents a mitochondrial-encoded or plastid-encoded protein. Enrichment in chloroplast samples (x-axis) or mitochondrial samples (y-axis) is calculated by dividing the PSM count for that protein from the respective purified organelles by the corresponding PSM count from total leaf samples. Point size is scaled based on total number of PSMs for that protein across all samples. Only proteins represented by at least 5 unique peptides in the dataset are shown, and enrichment/depletion values were capped at 10-fold for visualization purposes. The two biological replicates are averaged for this panel. In both panels, point shape indicates species identity (circle: *A. thaliana*; triangle: *S. conica*), and PSMs were excluded for peptides shared between multiple proteins.

**Figure S2.**
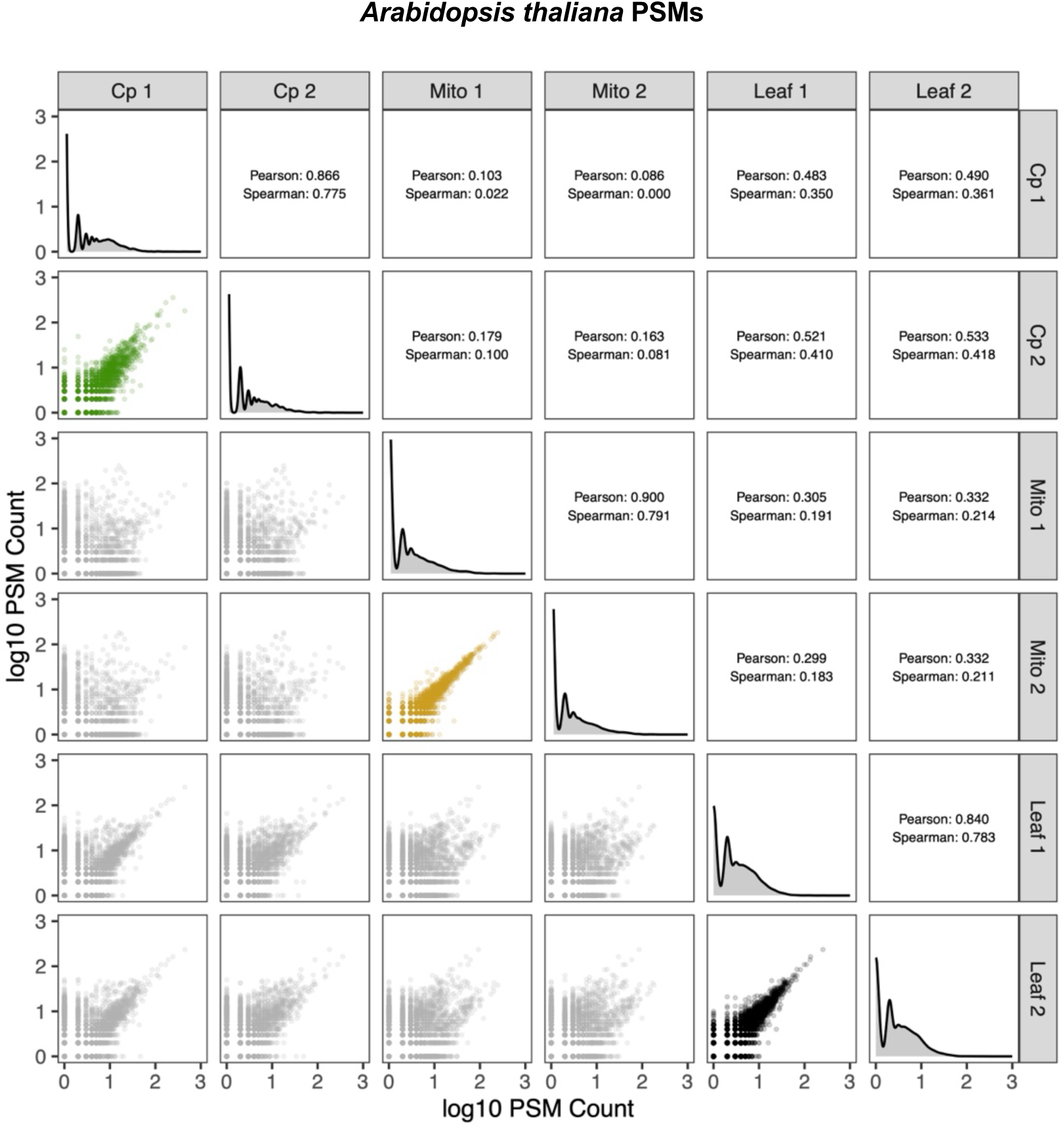
Pairwise correlation matrix among the six *A. thaliana* samples based on PSMs for each identified protein. Each point in the scatterplots below the downward diagonal represents a protein. The highlighted subplots show the three pairs of biological replicates of the same sample type that are expected to have high correlations. The correlation coefficients (Pearson’s *r* and Spearman’s *p*) corresponding to each subplot are shown above the downward diagonal. The downward diagonal itself shows the univariate distribution (density kernel) for PSM counts in each of the six samples. PSMs were excluded for peptides that were shared between multiple proteins. A baseline value of 1 was added to all values prior to log-transformation and visualization.

**Figure S3.**
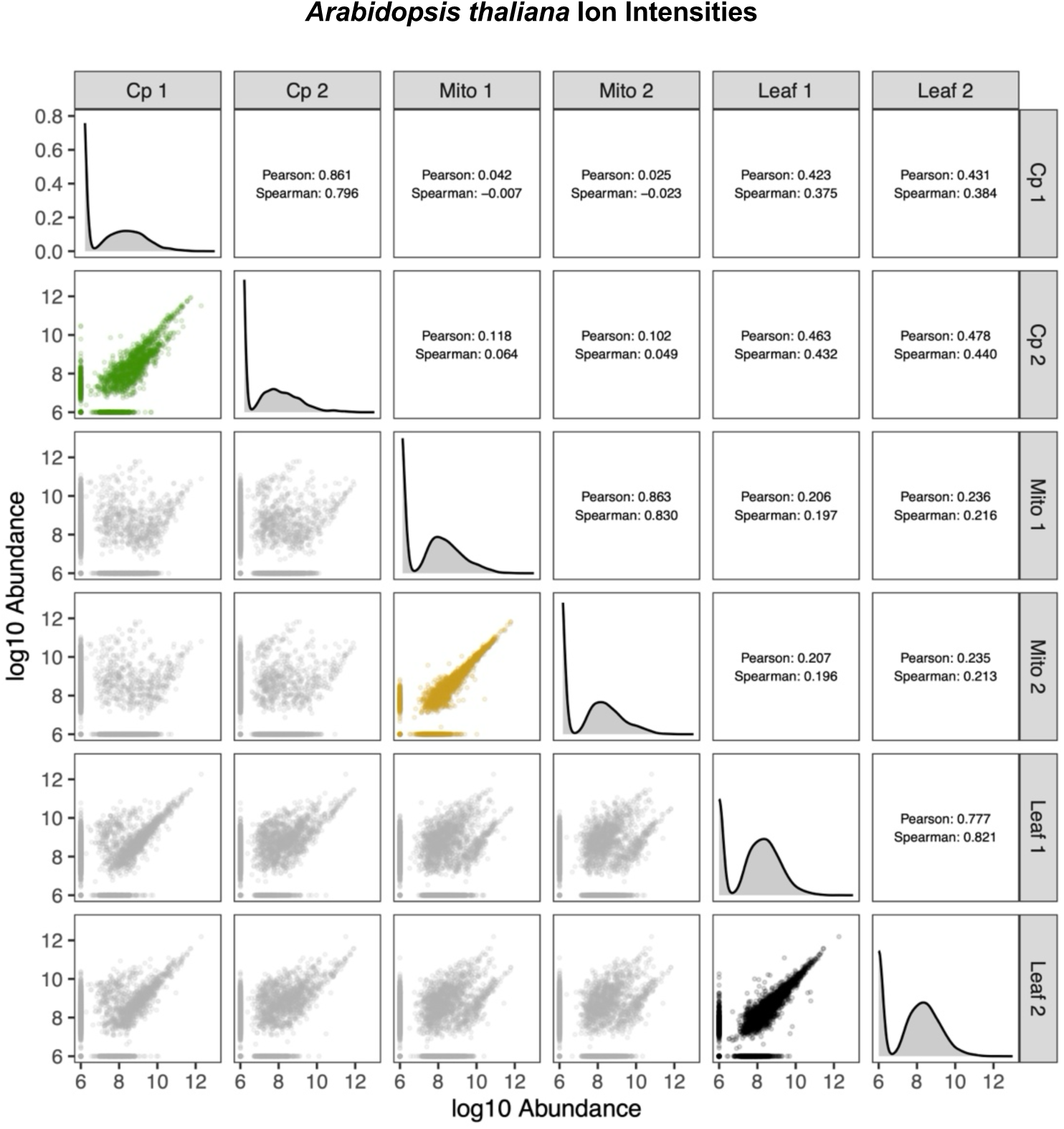
Pairwise correlation matrix among the six *A. thaliana* samples based on ion intensity for each identified protein. Each point in the scatterplots below the downward diagonal represents a protein. The highlighted subplots show the three pairs of biological replicates of the same sample type that are expected to have high correlations. The correlation coefficients (Pearson’s *r* and Spearman’s *p*) corresponding to each subplot are shown above the downward diagonal. The downward diagonal itself shows the univariate distribution (density kernel) for ion intensities in each of the six samples. Ion intensities were excluded for peptides that were shared between multiple proteins or that were identified solely based on an MS1 peak that was not validated in that sample with an MS2 spectrum. A baseline value of 1e6 was added to all values prior to log-transformation and visualization.

**Figure S4.**
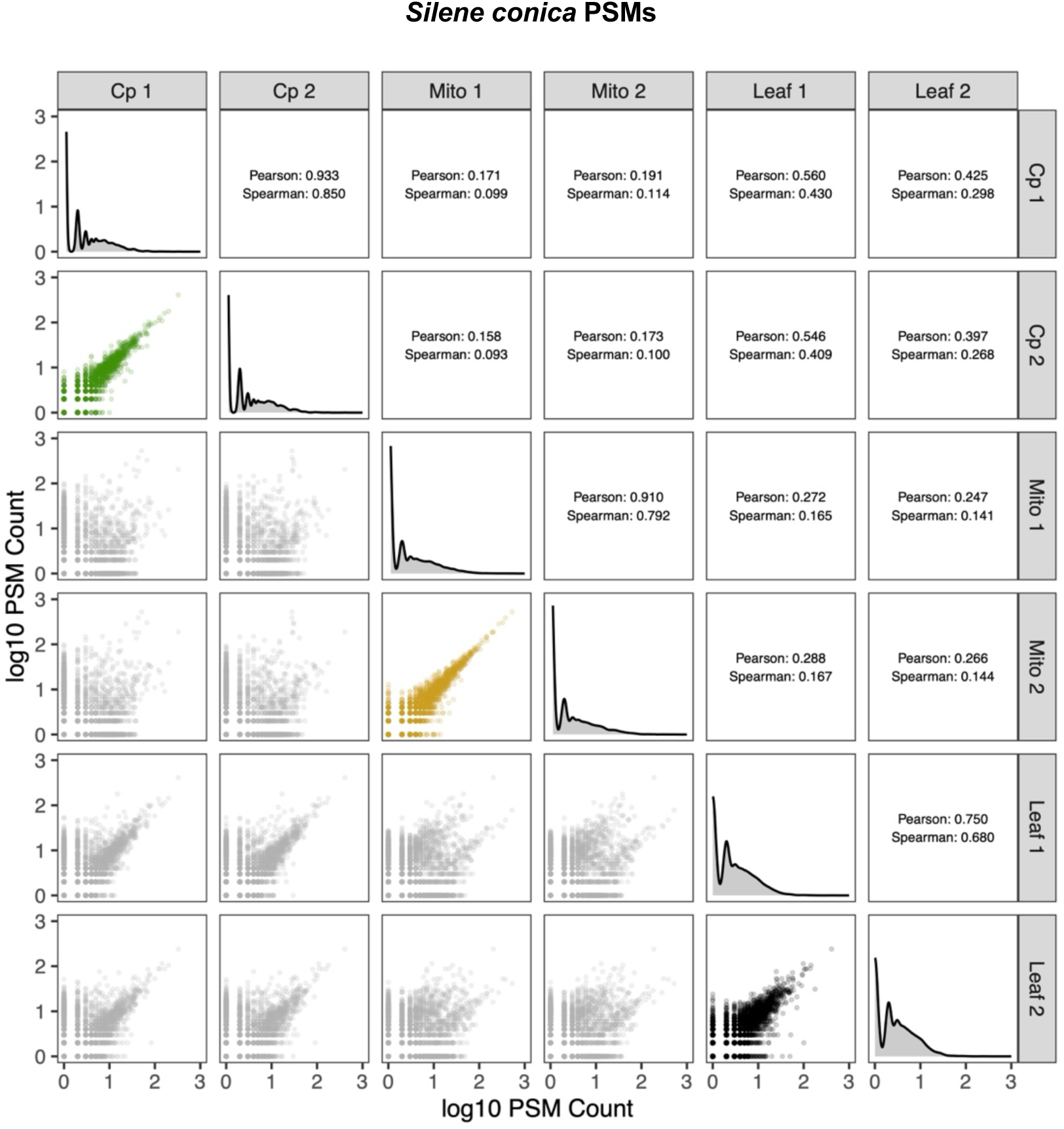
Pairwise correlation matrix among the six *S. conica* samples based on PSMs for each identified protein. Each point in the scatterplots below the downward diagonal represents a protein. The highlighted subplots show the three pairs of biological replicates of the same sample type that are expected to have high correlations. The correlation coefficients (Pearson’s *r* and Spearman’s *p*) corresponding to each subplot are shown above the downward diagonal. The downward diagonal itself shows the univariate distribution (density kernel) for PSM counts in each of the six samples. PSMs were excluded for peptides that were shared between multiple proteins. A baseline value of 1 was added to all values prior to log-transformation and visualization.

**Figure S5.**
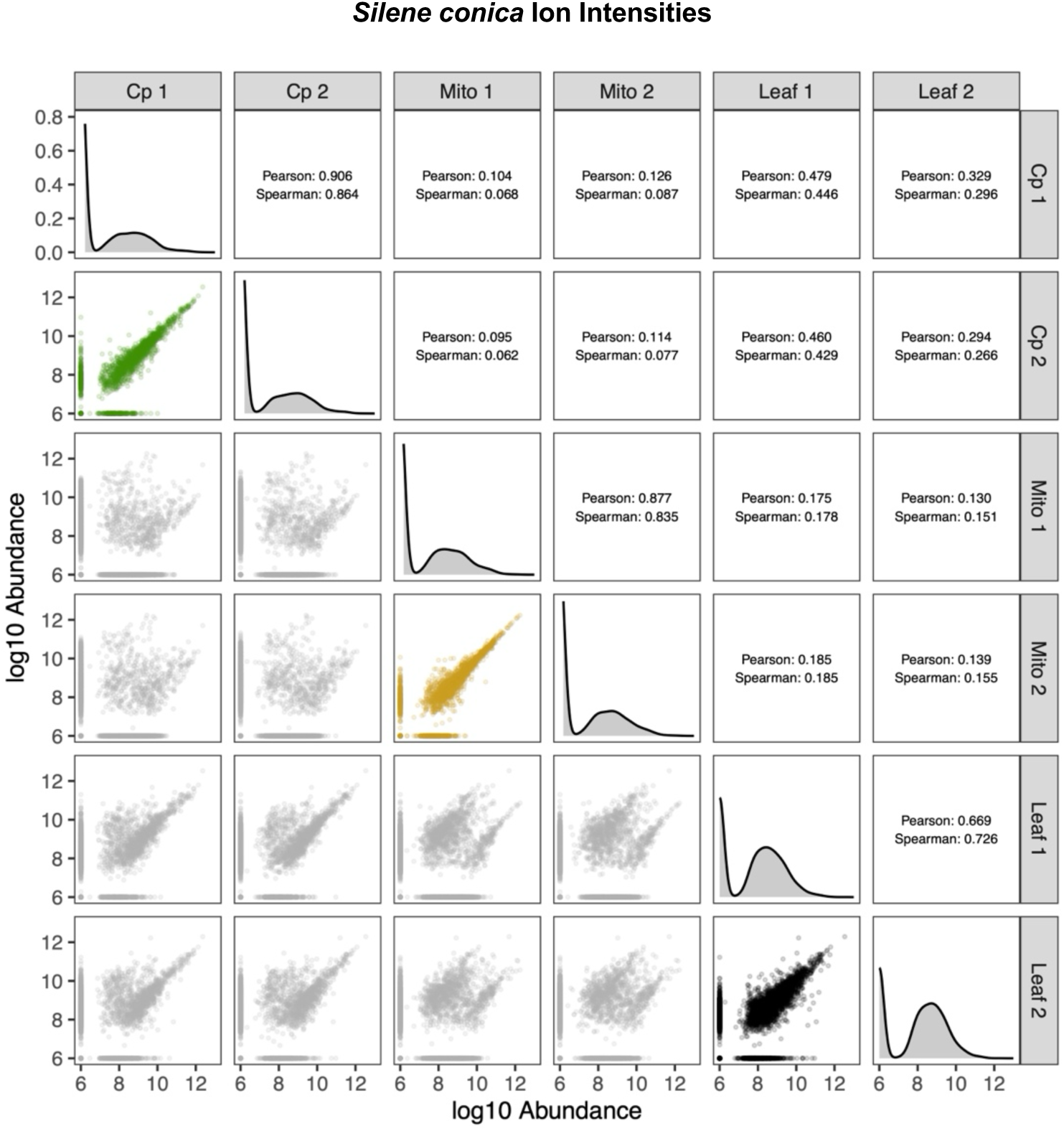
Pairwise correlation matrix among the six *S. conica* samples based on ion intensity for each identified protein. Each point in the scatterplots below the downward diagonal represents a protein. The highlighted subplots show the three pairs of biological replicates of the same sample type that are expected to have high correlations. The correlation coefficients (Pearson’s *r* and Spearman’s *p*) corresponding to each subplot are shown above the downward diagonal. The downward diagonal itself shows the univariate distribution (density kernel) for ion intensities in each of the six samples. Ion intensities were excluded for peptides that were shared between multiple proteins or that were identified solely based on an MS1 peak that was not validated in that sample with an MS2 spectrum. A baseline value of 1e6 was added to all values prior to log-transformation and visualization.

**Figure S6.**
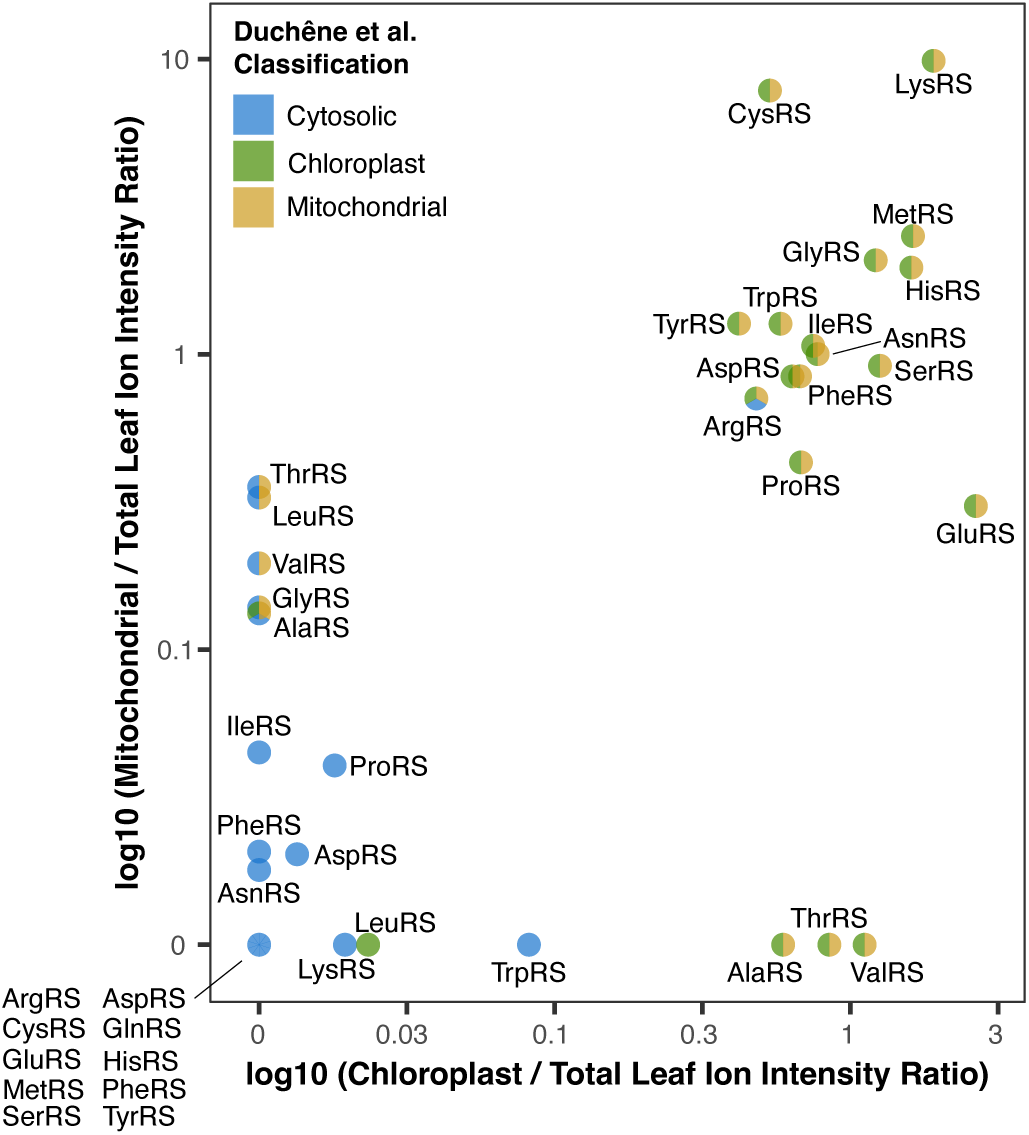
Summary of mitochondrial and chloroplast enrichment of *Arabidopsis thaliana* aaRSs relative to total leaf samples based on ratios of ion intensity averaged across two biological replicates. Ion intensities were excluded for peptides that were shared between multiple proteins or if they were identified solely based on an MS1 peak that was not validated in that sample with an MS2 spectrum. Color coding of points reflects whether the aaRS was previously classified as being targeted to the cytosol, chloroplasts, and/or mitochondria (Duchêne et al. 2005; Duchêne et al. 2009).

**Figure S7.**
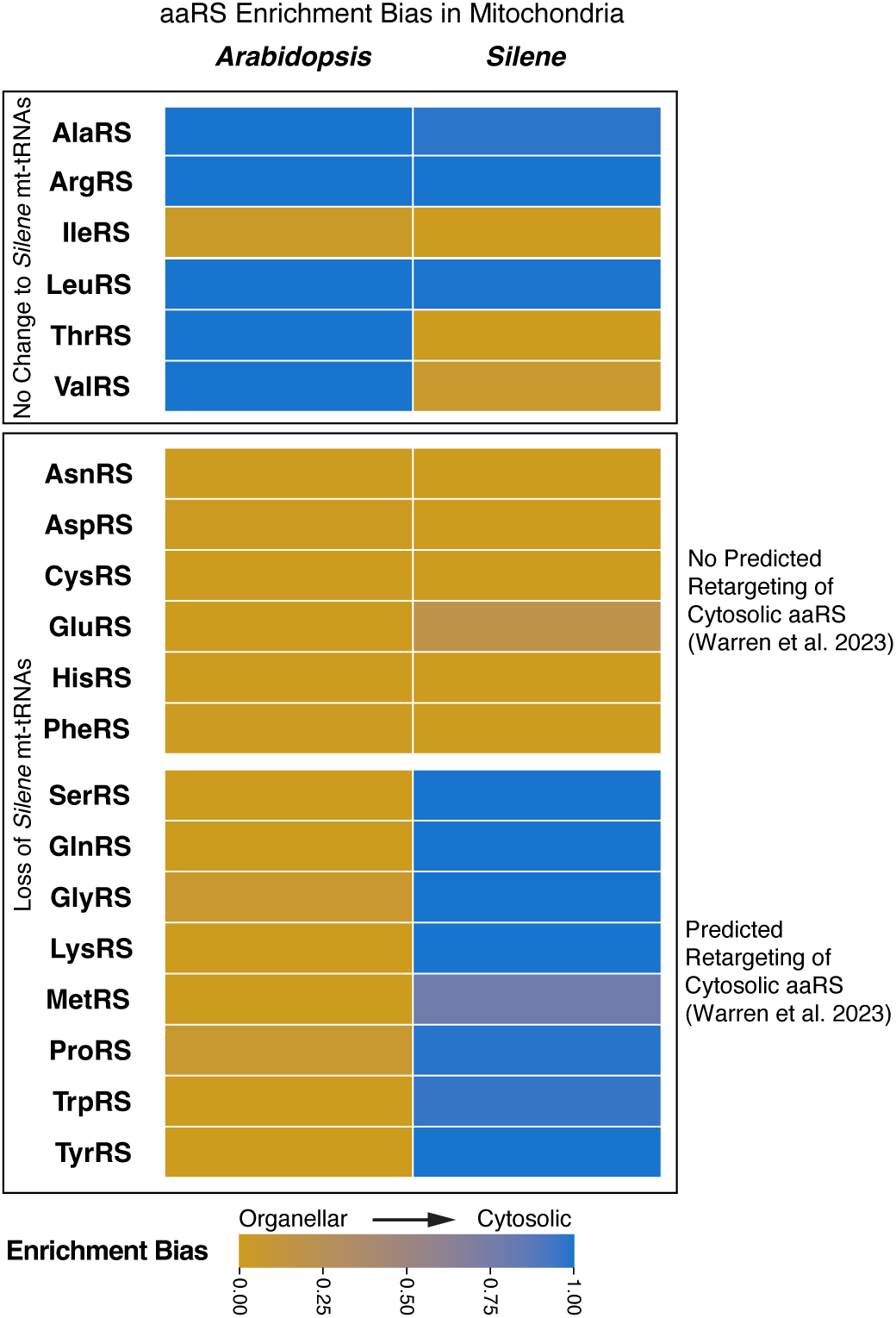
Summary of whether mitochondrial samples were biased towards containing organellar-like vs. cytosolic-like aaRSs based on proteomic analysis. This is the same representation as Figure 4 except that enrichment bias was calculated using ion intensity rather than PSM counts as the quantification metric. ion intensities were excluded for peptides that were shared between multiple proteins or if they were identified solely based on an MS1 peak that was not validated in that sample with an MS2 spectrum. See Methods and Figure 4 legend for more information.

**Figure S8.**
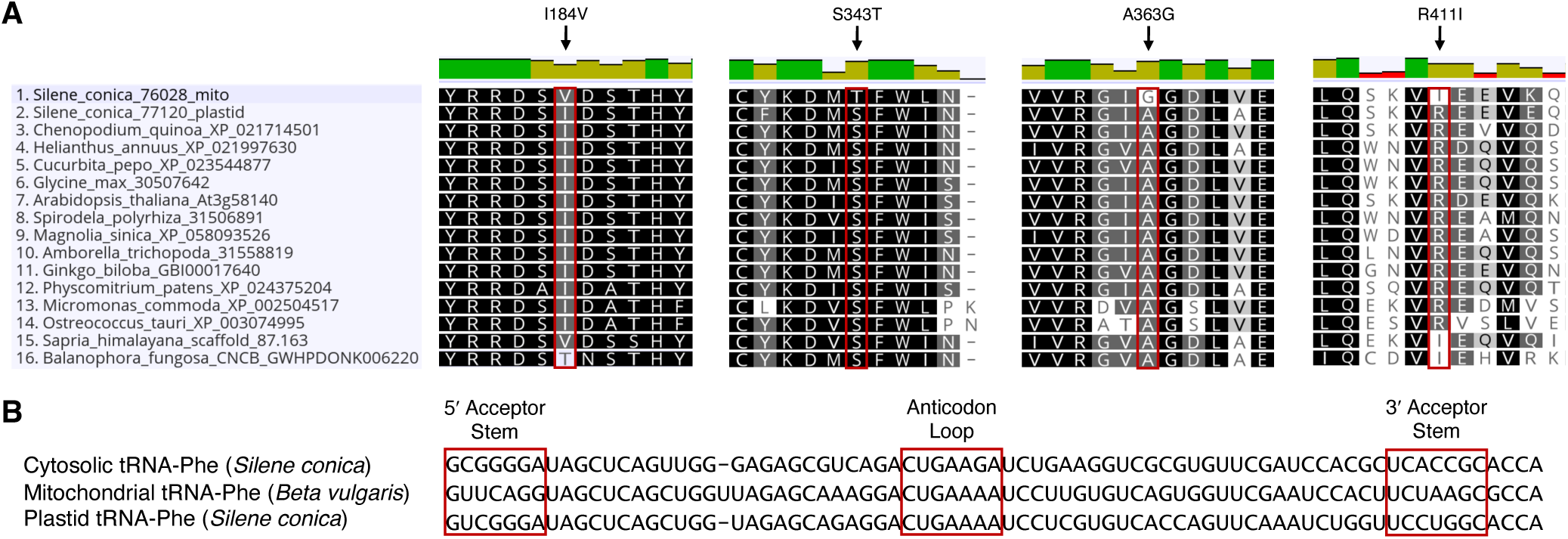
PheRS and tRNA-Phe alignments. (A) Amino acid alignments showing four positions in the PheRS sequence that have a derived change in the *Silene conica* mitochondrial PheRS but are otherwise conserved across a diverse sampling of green plants/algae. In two of these cases (I184V and R411I), parallel substitutions are observed in the parasitic plant taxa *Sapria himalayana* and *Balanophora fungosa*. (B) Alignment of cytosolic, mitochondrial, and plastid tRNA-Phe.

**Table S1.** Abundance of SUBA5-classified mitochondrial and plastid proteins in *A. thaliana* LC-MS/MS samples. The reported percentages reflect the total ion intensity or PSMs for proteins in the respective SUBA5 category divided by the total ion intensity or PSM signal for all identified proteins.

| Arabidopsis Sample | SUBA5 Mitochondrial Proteins |  |  | SUBA5 Plastid Proteins |  |  |
| --- | --- | --- | --- | --- | --- | --- |
|  | # Detected | Ion Intensity | PSMs | # Detected | Ion Intensity | PSMs |
| Mitochondrial 1 | 835 | 73.96% | 59.93% | 459 | 9.79% | 14.21% |
| Mitochondrial 2 | 783 | 73.16% | 61.25% | 363 | 10.15% | 13.27% |
| Leaf 1 | 459 | 6.79% | 11.29% | 1110 | 70.07% | 41.58% |
| Leaf 2 | 449 | 8.18% | 12.48% | 1090 | 69.65% | 43.07% |
| Chloroplast 1 | 176 | 2.06% | 5.25% | 1314 | 96.21% | 91.55% |
| Chloroplast 2 | 161 | 1.78% | 4.74% | 1138 | 95.54% | 86.56% |

**Table S2.** Normalized MS1 ion intensities for PheRS enzymes in *S. conica*.

| Sample | PheRS Type |  |  | Ratio of Mito to Chloro Types<sup>b</sup> |
| --- | --- | --- | --- | --- |
|  | Cytosolic<br>(Sconica_v3_11405-RA) | Chloroplast<br>(Sconica_v3_09891-RA) | Mitochondrial<br>(Chr01-anno1.g64625.t1) |  |
| Mt1 | 1.03e8 | 6.25e8 | 1.83e9 | 2.93 |
| Mt2 | 9.70e7 | 5.21e8 | 1.39e9 | 2.67 |
| Cp1 | 2.77e8 | 2.35e9 | 1.28e8 | 0.05 |
| Cp2 | 5.51e8 | 3.35e9 | 3.19e8 | 0.10 |
| Lf1 | 1.45e9 | 2.81e9 | 1.87e8 | 0.07 |
| Lf2 | 1.58e9 | 2.37e9 | 5.30e8 | 0.22 |

<sup>a</sup>Reported values exclude contributions from shared peptides but do include intensities reported only as “Peak Found” without PSM support. The identified differences would be even larger if these were excluded, as the putative mitochondrial PheRS produced no PSMs outside of the mitochondrial samples.
<sup>b</sup>The mitochondrial samples have a significantly higher ratio of mitochondrial to chloroplast PheRS types than the chloroplast samples (one-sided Student’s *t*-test of log-normalized ratios; *p* = 0.0029).

**Table S3.**
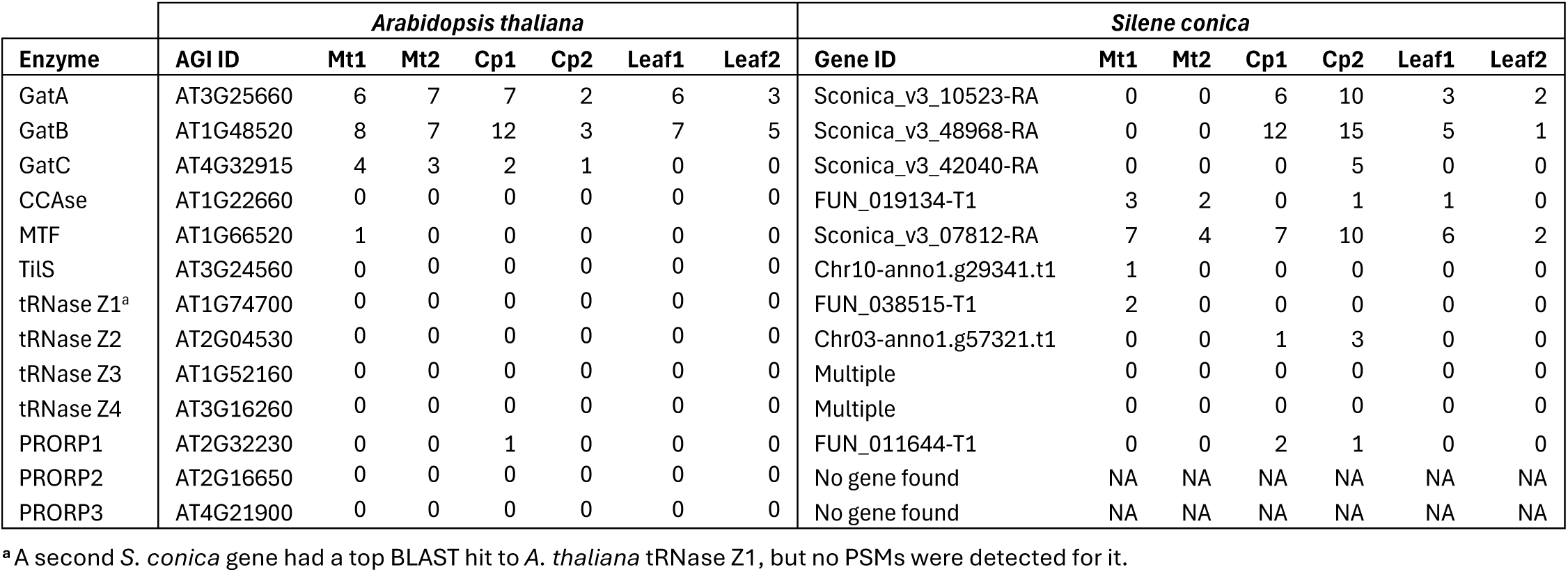
PSM counts by sample for additional enzymes involved in tRNA metabolism in mitochondria and other compartments.

**Table S4.**
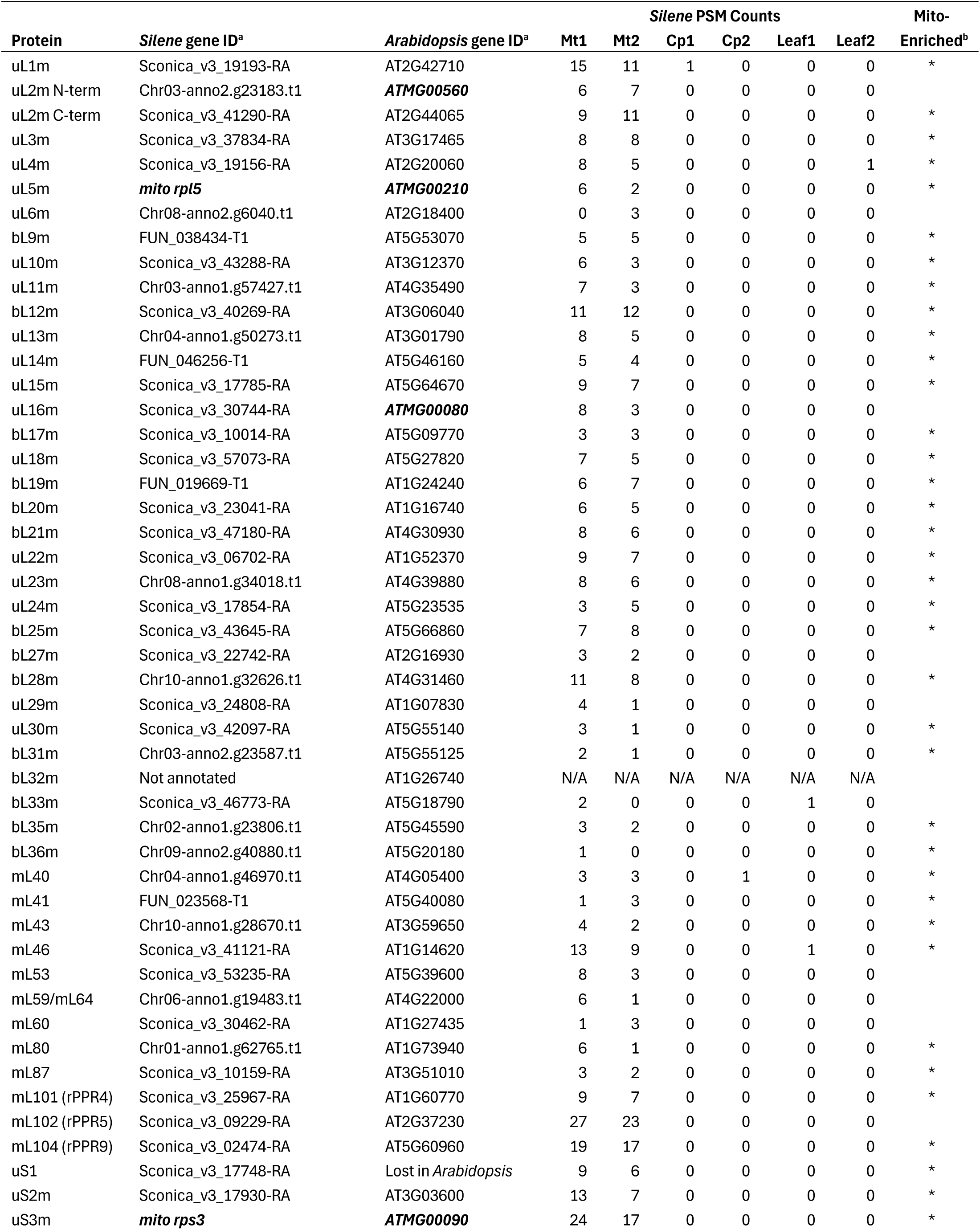
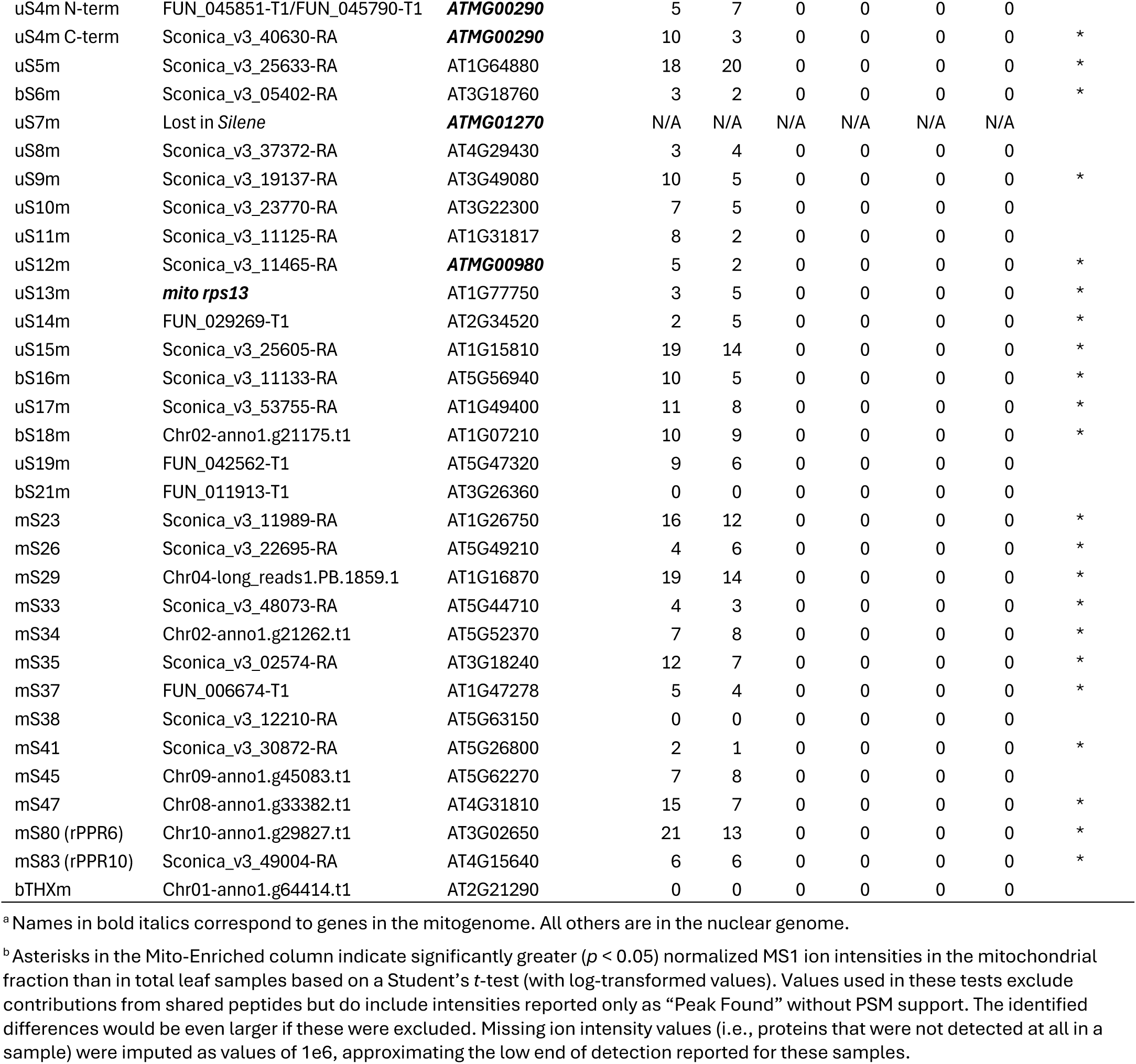
Genes encoding subunits of the *A. thaliana* mitoribosome (Waltz et al. 2020) and their counterparts in *S. conica*.

## References

Abramson J, Adler J, Dunger J, Evans R, Green T, Pritzel A, Ronneberger O, Willmore L, Ballard AJ, Bambrick J, et al. 2024. Accurate structure prediction of biomolecular interactions with AlphaFold 3. Nature 630:493–500.

Adams KL, Daley DO, Whelan J, Palmer JD. 2002. Genes for two mitochondrial ribosomal proteins in flowering plants are derived from their chloroplast or cytosolic counterparts. Plant Cell 14:931–943.

Adams KL, Ong HC, Palmer JD. 2001. Mitochondrial gene transfer in pieces: fission of the ribosomal protein gene rpl2 and partial or complete gene transfer to the nucleus. Mol. Biol. Evol. 18:2289–2297.

Adams KL, Palmer JD. 2003. Evolution of mitochondrial gene content: gene loss and transfer to the nucleus. Mol. Phylogenet. Evol. 29:380–395.

Adams KL, Qiu YL, Stoutemyer M, Palmer JD. 2002. Punctuated evolution of mitochondrial gene content: high and variable rates of mitochondrial gene loss and transfer to the nucleus during angiosperm evolution. Proc. Natl. Acad. Sci. U. S. A. 99:9905–9912.

Adams KL, Song K, Roessler PG, Nugent JM, Doyle JL, Doyle JJ, Palmer JD. 1999. Intracellular gene transfer in action: dual transcription and multiple silencings of nuclear and mitochondrial *cox2* genes in legumes. Proceedings of the National Academy of Sciences 96:13863–13868.

Berg M, Rogers R, Muralla R, Meinke D. 2005. Requirement of aminoacyl-tRNA synthetases for gametogenesis and embryo development in Arabidopsis: Aminoacyl-tRNA synthetase knockouts. Plant J. 44:866–878.

Berrissou C, Cognat V, Koechler S, Bergdoll M, Duchêne A-M, Drouard L. 2024. Extensive import of nucleus-encoded tRNAs into chloroplasts of the photosynthetic lycophyte, Selaginella kraussiana. Proc. Natl. Acad. Sci. U. S. A. 121:e2412221121.

Boore JL. 1999. Animal mitochondrial genomes. Nucleic Acids Res. 27:1767–1780.

von Braun SS, Sabetti A, Hanic-Joyce PJ, Gu J, Schleiff E, Joyce PBM. 2007. Dual targeting of the tRNA nucleotidyltransferase in plants: not just the signal. J. Exp. Bot. 58:4083–4093.

Broz AK, Waneka G, Wu Z, Sloan DB. 2021. Detecting de novo mitochondrial mutations in angiosperms with highly divergent evolutionary rates. Genetics 218:iyab039.

Canino G, Bocian E, Barbezier N, Echeverría M, Forner J, Binder S, Marchfelder A. 2009. Arabidopsis encodes four tRNase Z enzymes. Plant Physiol. 150:1494–1502.

Ceriotti LF, Gatica-Soria LM, Prasad KVSK, DeTar RA, Warren JM, Eichler E, Chustecki JM, Elowsky C, Christensen AC, Zhou R, et al. 2026. Reshaping organellar translation and tRNA metabolism: The consequences of photosynthesis loss and massive horizontal gene transfer. bioRxiv [Internet]:2026.01.09.698701. Available from: https://www.biorxiv.org/content/10.64898/2026.01.09.698701.abstract

Ceriotti LF, Roulet ME, Sanchez-Puerta MV. 2021. Plastomes in the holoparasitic family Balanophoraceae: Extremely high AT content, severe gene content reduction, and two independent genetic code changes. Mol. Phylogenet. Evol. 162:107208.

Chang Y-T, Barad BA, Rahmani H, Zid BM, Grotjahn DA. 2024. Cytoplasmic ribosomes on mitochondria alter the local membrane environment for protein import. bioRxivorg [Internet]. Available from: http://biorxiv.org/lookup/doi/10.1101/2024.07.17.604013

Cheng C-Y, Krishnakumar V, Chan AP, Thibaud-Nissen F, Schobel S, Town CD. 2017. Araport11: a complete reannotation of the Arabidopsis thaliana reference genome. Plant J. 89:789–804.

DeTar RA, Chustecki JM, Martinez-Hottovy A, Ceriotti LF, Broz AK, Lou X, Sanchez-Puerta MV, Elowsky C, Christensen AC, Sloan DB. 2024. Photosynthetic demands on translational machinery drive retention of redundant tRNA metabolism in plant organelles. Proc. Natl. Acad. Sci. U. S. A. 121:e2421485121.

Dimnet L, Salinas-Giegé T, Pullara S, Moyet L, Genevey C, Kuntz M, Duchêne A-M, Rolland N. 2024. Isolation of cytosolic ribosomes associated with plant mitochondria and chloroplasts. Methods Mol. Biol. 2776:289–302.

Duchêne A-M, Giritch A, Hoffmann B, Cognat V, Lancelin D, Peeters NM, Zaepfel M, Maréchal-Drouard L, Small ID. 2005. Dual targeting is the rule for organellar aminoacyl-tRNA synthetases in *Arabidopsis thaliana*. Proceedings of the National Academy of Sciences 102:16484–16489.

Duchêne AM, Peeters N, Dietrich A, Cosset A, Small ID, Wintz H. 2001. Overlapping destinations for two dual targeted glycyl-tRNA synthetases in Arabidopsis thaliana and Phaseolus vulgaris. J. Biol. Chem. 276:15275–15283.

Duchêne A-M, Pujol C, Maréchal-Drouard L. 2009. Import of tRNAs and aminoacyl-tRNA synthetases into mitochondria. Curr. Genet. 55:1–18.

Emms DM, Kelly S. 2022. SHOOT: phylogenetic gene search and ortholog inference. Genome Biol. 23:85.

Fields P, Weber MM, Waneka G, Broz A, Sloan DB. 2023. Chromosome-level genome assembly for the angiosperm Silene conica. Genome Biol. Evol. 15:evad192.

Fields PD, Waneka G, Naish M, Schatz MC, Henderson IR, Sloan DB. 2022. Complete Sequence of a 641-kb Insertion of Mitochondrial DNA in the Arabidopsis thaliana Nuclear Genome. Genome Biol. Evol. 14:evac059.

Fu L, Niu B, Zhu Z, Wu S, Li W. 2012. CD-HIT: accelerated for clustering the next-generation sequencing data. Bioinformatics 28:3150–3152.

Fuchs P, Rugen N, Carrie C, Elsässer M, Finkemeier I, Giese J, Hildebrandt TM, Kühn K, Maurino VG, Ruberti C. 2020. Single organelle function and organization as estimated from Arabidopsis mitochondrial proteomics. Plant J. 101:420–441.

Gamper H, Hou Y-M. 2020. A label-free assay for aminoacylation of tRNA. Genes 11:1173.

Gobert A, Gutmann B, Taschner A, Gössringer M, Holzmann J, Hartmann RK, Rossmanith W, Giegé P. 2010. A single Arabidopsis organellar protein has RNase P activity. Nat. Struct. Mol. Biol. 17:740–744.

Gould SB, Waller RF, McFadden GI. 2008. Plastid evolution. Annu. Rev. Plant Biol. 59:491–517.

Gutmann B, Gobert A, Giegé P. 2012. PRORP proteins support RNase P activity in both organelles and the nucleus in Arabidopsis. Genes Dev. 26:1022–1027.

Heinemann B, Künzler P, Eubel H, Braun H-P, Hildebrandt TM. 2021. Estimating the number of protein molecules in a plant cell: protein and amino acid homeostasis during drought. Plant Physiol. 185:385–404.

Hooper CM, Tanz SK, Castleden IR, Vacher MA, Small ID, Millar AH. 2014. SUBAcon: a consensus algorithm for unifying the subcellular localization data of the Arabidopsis proteome. Bioinformatics 30:3356–3364.

Ibba M, Söll D. 2004. Aminoacyl-tRNAs: setting the limits of the genetic code. Genes Dev. 18:731–738.

Katoh K, Standley DM. 2013. MAFFT multiple sequence alignment software version 7: improvements in performance and usability. Mol. Biol. Evol. 30:772–780.

Kearse M, Moir R, Wilson A, Stones-Havas S, Cheung M, Sturrock S, Buxton S, Cooper A, Markowitz S, Duran C. 2012. Geneious Basic: an integrated and extendable desktop software platform for the organization and analysis of sequence data. Bioinformatics 28:1647–1649.

Kley J, Heil M, Muck A, Svatos A, Boland W. 2010. Isolating intact chloroplasts from small Arabidopsis samples for proteomic studies. Anal. Biochem. 398:198–202.

Klipcan L, Moor N, Finarov I, Kessler N, Sukhanova M, Safro MG. 2012. Crystal structure of human mitochondrial PheRS complexed with tRNAPhe in the active “open” state. J. Mol. Biol. 415:527–537.

Kubo N, Arimura S. 2010. Discovery of the rpl10 gene in diverse plant mitochondrial genomes and its probable replacement by the nuclear gene for chloroplast RPL10 in two lineages of angiosperms. DNA Res. 17:1–9.

Liu SL, Zhuang Y, Zhang P, Adams KL. 2009. Comparative analysis of structural diversity and sequence evolution in plant mitochondrial genes transferred to the nucleus. Mol. Biol. Evol. 26:875–891.

McCutcheon JP, Garber AI, Spencer N, Warren JM. 2024. How do bacterial endosymbionts work with so few genes? PLoS Biol. 22:e3002577.

McCutcheon JP, Moran NA. 2012. Extreme genome reduction in symbiotic bacteria. Nat. Rev. Microbiol. 10:13–26.

Meiklejohn CD, Holmbeck MA, Siddiq MA, Abt DN, Rand DM, Montooth KL. 2013. An Incompatibility between a mitochondrial tRNA and its nuclear-encoded tRNA synthetase compromises development and fitness in Drosophila. PLoS Genet. 9:e1003238.

Meyer EH, Tomaz T, Carroll AJ, Estavillo G, Delannoy E, Tanz SK, Small ID, Pogson BJ, Millar AH. 2009. Remodeled respiration in ndufs4 with low phosphorylation efficiency suppresses Arabidopsis germination and growth and alters control of metabolism at night. Plant Physiol. 151:603–619.

Mian S, Leitch IJ. 2024. The genome sequence of the corn cockle, Agrostemma githago L., 1753 (Caryophyllaceae). Wellcome Open Res. 9:590.

Mireau H, Lancelin D, Small ID. 1996. The same Arabidopsis gene encodes both cytosolic and mitochondrial alanyl-tRNA synthetases. Plant Cell 8:1027–1039.

Molina J, Hazzouri KM, Nickrent D, Geisler M, Meyer RS, Pentony MM, Flowers JM, Pelser P, Barcelona J, Inovejas SA. 2014. Possible loss of the chloroplast genome in the parasitic flowering plant *Rafflesia lagascae* (Rafflesiaceae). Mol. Biol. Evol. 31:793–803.

Mower JP, Bonen L. 2009. Ribosomal protein L10 is encoded in the mitochondrial genome of many land plants and green algae. BMC Evol. Biol. 9:265.

Peeters NM, Chapron A, Giritch A, Grandjean O, Lancelin D, Lhomme T, Vivrel A, Small I. 2000. Duplication and quadruplication of Arabidopsis thaliana cysteinyl- and asparaginyl-tRNA synthetase genes of organellar origin. J. Mol. Evol. 50:413–423.

Pérez-Riverol Y, Bandla C, Kundu DJ, Kamatchinathan S, Bai J, Hewapathirana S, John NS, Prakash A, Walzer M, Wang S, et al. 2024. The PRIDE database at 20 years: 2025 update. Nucleic Acids Res. 53:D543–D553.

Pett W, Lavrov DV. 2015. Cytonuclear interactions in the evolution of animal mitochondrial tRNA metabolism. Genome Biol. Evol. 7:2089–2101.

Pruitt K, Tatusova T, Maglott D. 2004. NCBI Reference Sequence (RefSeq): a curated non-redundant sequence database of genomes, transcripts and proteins. Nucleic Acids Research 33:D501–D504.

Pujol C, Bailly M, Kern D, Marechal-Drouard L, Becker H, Duchene AM. 2008. Dual-targeted tRNA-dependent amidotransferase ensures both mitochondrial and chloroplastic Gln-tRNAGln synthesis in plants. Proceedings of the National Academy of Sciences 105:6481–6485.

Richardson AO, Rice DW, Young GJ, Alverson AJ, Palmer JD. 2013. The “fossilized” mitochondrial genome of Liriodendron tulipifera: ancestral gene content and order, ancestral editing sites, and extraordinarily low mutation rate. BMC Biol. 11:29.

Roger AJ, Muñoz-Gómez SA, Kamikawa R. 2017. The origin and diversification of mitochondria. Curr. Biol. 27:R1177–R1192.

Rogers KC, Söll D. 1995. Divergence of glutamate and glutamine aminoacylation pathways: providing the evolutionary rationale for mischarging. J. Mol. Evol. 40:476–481.

Rugen N, Senkler M, Braun H-P. 2024. Deep proteomics reveals incorporation of unedited proteins into mitochondrial protein complexes in Arabidopsis. Plant Physiol. 195:1180–1199.

Rugen N, Straube H, Franken LE, Braun H-P, Eubel H. 2019. Complexome profiling reveals association of PPR proteins with ribosomes in the mitochondria of plants. Mol. Cell. Proteomics 18:1345–1362.

Salinas-Giegé T, Giegé R, Giegé P. 2015. tRNA biology in mitochondria. Int. J. Mol. Sci. 16:4518–4559.

Skaltsogiannis V, Nguyen T-T, Corre N, Pflieger D, Blevins T, Hashem Y, Giegé P, Waltz F. 2024. Structural insights into maturation and translation of a plant mitoribosome. bioRxiv [Internet]. Available from: 10.1101/2024.10.28.620559

Skippington E, Barkman TJ, Rice DW, Palmer JD. 2015. Miniaturized mitogenome of the parasitic plant Viscum scurruloideum is extremely divergent and dynamic and has lost all nad genes. Proceedings of the National Academy of Sciences [Internet] In Press. Available from: 10.1073/pnas.1504491112 [doi]

Sloan DB, Alverson AJ, Chuckalovcak JP, Wu M, McCauley DE, Palmer JD, Taylor DR. 2012. Rapid evolution of enormous, multichromosomal genomes in flowering plant mitochondria with exceptionally high mutation rates. PLoS Biol. 10:e1001241.

Sloan DB, Alverson AJ, Wu M, Palmer JD, Taylor DR. 2012. Recent acceleration of plastid sequence and structural evolution coincides with extreme mitochondrial divergence in the angiosperm genus Silene. Genome Biol. Evol. 4:294–306.

Sloan DB, Triant DA, Forrester NJ, Bergner LM, Wu M, Taylor DR. 2014. A recurring syndrome of accelerated plastid genome evolution in the angiosperm tribe Sileneae (Caryophyllaceae). Mol. Phylogenet. Evol. 72:82–89.

Sloan DB, Warren JM, Williams AM, Kuster SA, Forsythe ES. 2023. Incompatibility and interchangeability in molecular evolution. Genome Biol. Evol. 15:evac184.

Sloan DB, Warren JM, Williams AM, Wu Z, Abdel-Ghany SE, Chicco AJ, Havird JC. 2018. Cytonuclear integration and co-evolution. Nat. Rev. Genet. 19:635–648.

Smith D, Asmail S. 2014. Next-generation sequencing data suggest that certain nonphotosynthetic green plants have lost their plastid genomes. New Phytol. 204:7–11.

Souciet G, Menand B, Ovesna J, Cosset A, Dietrich A, Wintz H. 1999. Characterization of two bifunctional Arabdopsis thaliana genes coding for mitochondrial and cytosolic forms of valyl-tRNA synthetase and threonyl-tRNA synthetase by alternative use of two in-frame AUGs: A. thalianagene characterization using two in-frame AUGs. Eur. J. Biochem. 266:848–854.

Steinmetz A, Weil J. 1986. Isolation and characterization of chloroplast and cytoplasmic transfer RNAs. Methods in Enzymology 118:212–231.

Stupar RM, Lilly JW, Town CD, Cheng Z, Kaul S, Buell CR, Jiang J. 2001. Complex mtDNA constitutes an approximate 620-kb insertion on Arabidopsis thaliana chromosome 2: implication of potential sequencing errors caused by large-unit repeats. Proc. Natl. Acad. Sci. U. S. A. 98:5099–5103.

Su H-J, Barkman TJ, Hao W, Jones SS, Naumann J, Skippington E, Wafula EK, Hu J-M, Palmer JD, DePamphilis CW. 2019. Novel genetic code and record-setting AT-richness in the highly reduced plastid genome of the holoparasitic plant Balanophora. Proceedings of the National Academy of Sciences 116:934–943.

Suzuki T, Miyauchi K. 2010. Discovery and characterization of tRNAIle lysidine synthetase (TilS). FEBS Lett. 584:272–277.

Timmis JN, Ayliffe MA, Huang CY, Martin W. 2004. Endosymbiotic gene transfer: Organelle genomes forge eukaryotic chromosomes. Nat. Rev. Genet. 5:123–135.

Uwer U, Willmitzer L, Altmann T. 1998. Inactivation of a glycyl-tRNA synthetase leads to an arrest in plant embryo development. Plant Cell 10:1277–1294.

Waltz F, Nguyen TT, Arrivé M, Bochler A, Chicher J, Hammann P, Kuhn L, Quadrado M, Mireau H, Hashem Y. 2019. Small is big in Arabidopsis mitochondrial ribosome. Nature Plants 5:106–117.

Waltz F, Soufari H, Bochler A, Giegé P, Hashem Y. 2020. Cryo-EM structure of the RNA-rich plant mitochondrial ribosome. Nat. Plants 6:377–383.

Warren JM, Broz AK, Martinez-Hottovy A, Elowsky C, Christensen AC, Sloan DB. 2023. Rewiring of aminoacyl-tRNA synthetase localization and interactions in plants with extensive mitochondrial tRNA gene loss. Mol. Biol. Evol. 40:msad163.

Warren JM, Salinas-Giegé T, Triant DA, Taylor DR, Drouard L, Sloan DB. 2021. Rapid shifts in mitochondrial tRNA import in a plant lineage with extensive mitochondrial tRNA gene loss. Mol. Biol. Evol. 38:5735–5751.

Warren JM, Sloan DB. 2020. Interchangeable parts: The evolutionarily dynamic tRNA population in plant mitochondria. Mitochondrion 52:144–156.

van Wijk KJ, Bentolila S, Leppert T, Sun Q, Sun Z, Mendoza L, Li M, Deutsch EW. 2024. Detection and editing of the updated Arabidopsis plastid- and mitochondrial-encoded proteomes through PeptideAtlas. Plant Physiol. 194:1411–1430.

van Wijk KJ, Leppert T, Sun Q, Boguraev SS, Sun Z, Mendoza L, Deutsch EW. 2021. The Arabidopsis PeptideAtlas: Harnessing worldwide proteomics data to create a comprehensive community proteomics resource. Plant Cell 33:3421–3453.

van Wijk KJ, Peltier JB, Giacomelli L. 2007. Isolation of chloroplast proteins from *Arabidopsis thaliana* for proteome analysis. Methods Mol. Biol. 355:43–48.

Yu R, Zhi X, Ceriotti LF, Skippington E, Rice DW, Su H-J, Barkman TJ, Sun C, Liu Y, Fang D, et al. 2025. A record-setting mitogenome in the holoparasitic plant Balanophora yakushimensis accompanied by exceptional loss of organellar DNA repair and recombination genes. BMC Biol [Internet] 23. Available from: 10.1186/s12915-025-02449-8

